# Modelling the impact of magnesium ions concentration on the folding of the SAM-II riboswitch

**DOI:** 10.1101/2021.04.12.439486

**Authors:** Osama Alaidi

## Abstract

Riboswitches are regulatory elements present in bacterial messenger RNA acting as sensors of small molecules and consequently playing a vital role in bacterial gene regulation. The SAM-II riboswitch is a class of riboswitches, that recognizes S-adenosyl methionine. It has been previously illustrated that the presence of Mg^2+^ ions stabilizes the pre-existing minor state of the riboswitch, which is structurally characterised by having a nucleated pseudoknot, leading to the increase of its probability. In this study, an analytical equilibrium model is developed to describe the impact of Mg^2+^ ions concentration on the folding of the SAM-II riboswitch, linking RNA folding and tertiary interactions energetics to ligand binding, and, hence enabling quantitative predictions. The method was used to study the role of the P1 helix sequence in determining the fraction of binding competent conformers of the SAM-II riboswitch, by simulating the Mg^2+^ titration curves of various mutants.

**Graphical abstract:** 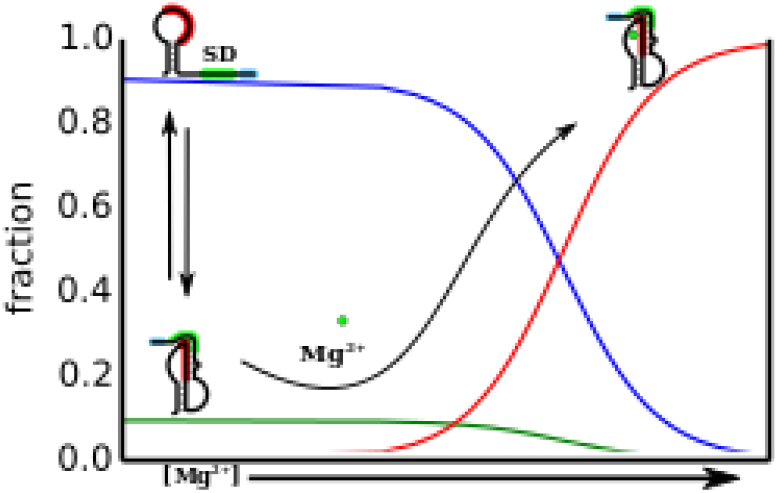

## 1 INTRODUCTION

Magnesium-induced RNA tertiary interactions are widespread [1–6]. Numerous studies show that various classes of RNA undergo specific structural changes, when placed in an environment having concentrations of magnesium ions comparable to cellular concentrations. Many of the latter structural changes are functionally important tertiary interactions [2–4,7,8]. In riboswitches, tertiary interactions induced by magnesium ions, usually influence the riboswitch binding with its cognate ligand [7,9,10]. The number of magnesium ions that interact with the RNA and the affinity of the RNA to these ions are variable [5,11]. Crystal structures of riboswitches often show one or more magnesium ions bound to the RNA [2,11–13] though in some cases these ions can be found in unexpected locations or even absent [14,15] despite the clear evidence that these riboswitches undergo structural changes in the presence of these ions [7,15].

The SAM-II riboswitch (**Figure 1**) is an example of such riboswitches that are influenced by Mg^2+^ ions. The riboswitch forms a set of tertiary interactions involving the formation of a classic H-type pseudoknot. The subsequent binding of its cognate ligand, S-adenosyl methionine (SAM), results in the sequestration of the ribosome binding site (Shine-Dalgarno (SD) sequence) [14,16], and eventually translation termination [16]. The folding of the riboswitch due to the pseudoknot formation, along with other tertiary interactions, leads to RNA compaction [7,17]. The crystal structure of the riboswitch shows that the ligand is partially buried under the surface (**Figure 1B**), suggesting that ligand binding is accompanied by subsequent conformational changes. Consequently, several studies have suggested a conformational capture mechanism in which SAM recognises the conformers that possesses the pre-nucleated pseudoknot [3,7,18–20].

**Figure 1.**
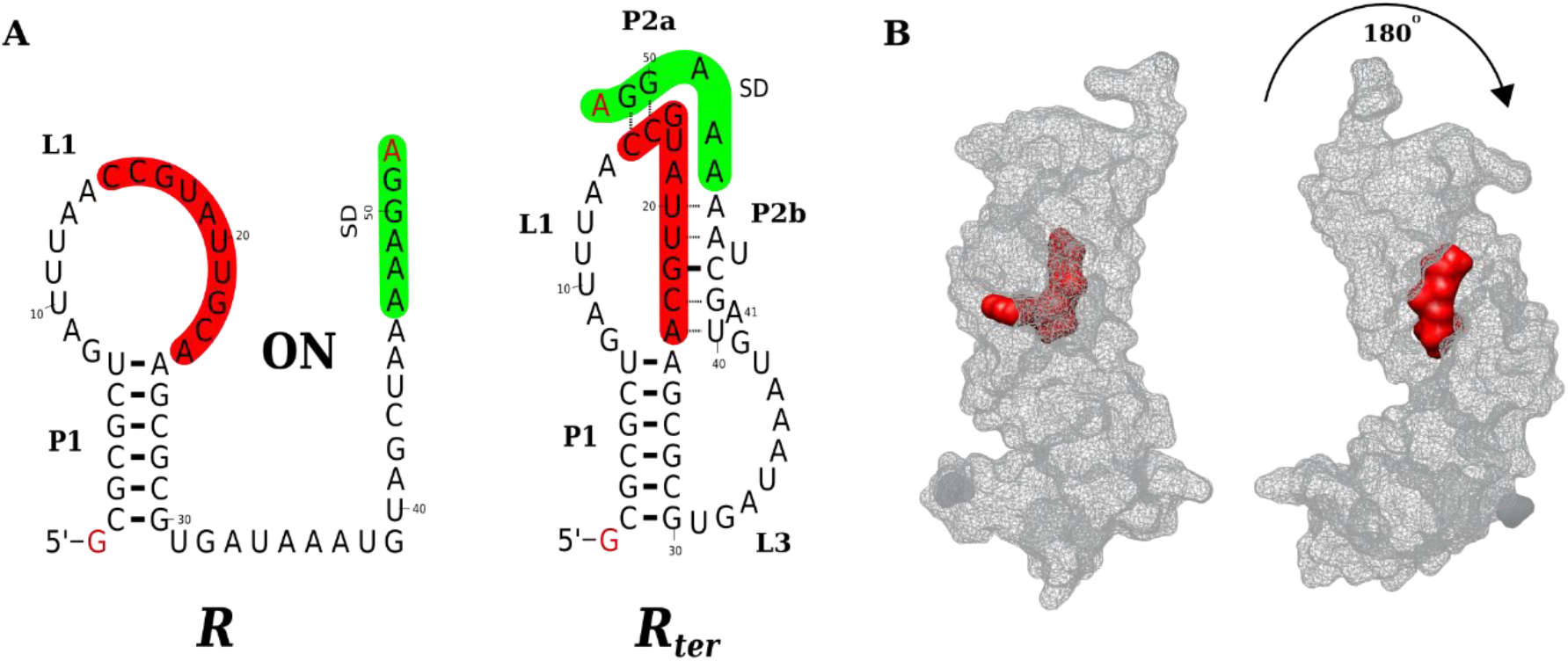
The structure of the SAM-II Riboswitch. (**A**) The figure illustrates the secondary structure of the Sargasso Sea SAM-II riboswitch upstream *metX* gene, illustrating the expected base pairing of the experimental sequence. The first state (named *R*) represent the ON state. In the latter, a pseudoknot is not formed and hence the SD region is available for ribosome binding. The SD and the anti-SD regions are shaded in green and red, respectively. The second conformer, *R*_*ter*_ is an intermediate with a nucleated pseudoknot, that is stabilized by the binding of Mg^2+^ ions. The latter conformer, which is used within the model developed in this study, is based on experiments that probed the interaction of the base pairs G22 and C43 (base pair interaction is represented by a solid line), indicating the nucleation of the pseudoknot [3]. Indeed, other pseudoknot Watson and Crick (WC) base pairs (interactions are represented by dashed lines) as well as non-WC base pairing and interactions (omitted from here for simplicity) may or may not be formed. (**B**) The figure shows different views of the wire surface representation of the SAM-II riboswitch and its cognate ligand, SAM, (in grey and red colors, respectively). Clearly, the ligand is partially buried under the surface of the RNA, suggesting that structural changes must have occurred, after the binding of the ligand to the competent conformers. The figure was produced from the crystal structure (PDB ID *2qwy*) [14].

The impact of magnesium ions on the SAM-II riboswitch have been a subject of several studies, in which controversial mechanisms have been proposed. The crystal structure illustrated the presence of a single magnesium ion that is far away from the SAM binding site, located near the phosphate groups of the nucleotides at 5’ termini, positioned between two RNA molecules [14]. Several subsequent studies had concluded that multiple conformations exist in solution [18] and that the presence of magnesium ions in the solvent promotes tertiary interactions that lead to the nucleation of the pseudoknot [3,7,19]. Based on NMR data, it has been suggested that magnesium ions most likely induce the formation of new tertiary loop-loop or loop-stem interactions, as evident from the heavy perturbation of the imino proton of a residue in the L1 loop (U12). Such interactions are likely to result in the pseudoknot nucleation and consequently the compaction of the unfolded state of the riboswitch [7]. This observation suggested a specific Mg^2+^ binding site near the SAM binding pocket, where it induces its major effect. On the contrary, a later computational study suggested that such structural changes can be explained by the non-specific interactions of the magnesium ions, present in the solvent layer around the RNA, with the backbone phosphate groups [17]. The latter conclusion was mainly based on comparing changes in the radius of gyration of the RNA during the simulation, with those seen experimentally in size exclusion chromatography (SEC) and small angle X-ray scattering (SAXS).

More recently however, the crystal structure of the SAM-V riboswitch [11,21], which is also a H-type pseudoknot and have a similar structure to the SAM-II riboswitch (**Figure S1**), showed that a hydrated metal ion (interpreted as a magnesium ion) in the L1 loop, near the ligand binding site, is present. The Mg^2+^ ion was found to be bound to the U10 phosphate oxygen, three water molecules and the SAM carboxyl group [11], effectively aiding in the formation of the triplex and further stabilizing the binding of the cognate ligand. This site is very similar to the site that had been described earlier by Chen et al [7] and observed to be heavily perturbed by Mg^2+^ ions, strongly suggesting that the binding of magnesium to a specific site or residue is likely to play a crucial role in the formation of the observed structural changes and its associated tertiary interactions in the SAM-II riboswitch, similar to that were seen in the SAM-V riboswitch. The latter interactions eventually would lead to the compaction of the RNA.

Motivated by such findings, in this study, a simple conformational capture model is used to quantitively predict the impact of magnesium ions on the SAM-II riboswitch. The simple analytical model is shown to be able to predict experimental observables at equilibrium. Despite that the model is developed to describe the influence of magnesium ions on the SAM-II riboswitch from Sargasso Sea metagenome [22], it can be generalized and adapted to other RNA where a structural change is induced by a ligand.

## 2 METHODS

### 2.1 The development of a conformational capture model

The secondary structures of the SAM-II riboswitch from the Sargasso Sea metagenome [22] is illustrated in **Figure 1A**. The ON and OFF signals relay on the formation of a pseudoknot that leads to sequestering the ribosome binding site, i.e. Shine-Dalgarno (SD) region. The first step in the formation of this pseudoknot is likely to be a nucleation process that involves the formation of one or more base pairs. Multiple studies have suggested that conformers with a nucleated or partially formed pseudoknot are sampled in the absence of SAM, and that these conformers are likely to be SAM binding competent, proposing a conformational capture mechanism [3,7,18,19]. Thus, to be able to accurately model SAM binding and its associated structural changes, one should consider quantitively defining the fraction that possesses such nucleated or partially formed pseudoknot.

A conformational capture mechanism implies that conformers having specific structural criteria, that enables a complete pseudoknot to be formed, are considered *binding competent* conformers for both Mg^2+^ and SAM. Conformers lacking such structural features are considered as *binding incompetent*. Indeed, conformers that have a nucleated or partially formed pseudoknot would enable the formation of the complete pseudoknot and triplex upon SAM binding. For this nucleation to occur in a conformer, certain secondary structure elements or criteria need to be present. Since the presence of the latter elements will determine the nucleation of the tertiary interactions, they would also, eventually, determine Mg^2+^ and SAM binding. Using this logic and proposed folding steps a model was constructed. The schematic diagram in **Figure 2** illustrates models for the RNA folding steps in the absence, panel (**A**), and in the presence, panel (**B**), of Mg^2+^ ions. The RNA species involved in this model are defined as follows; *R*_0_ is the concentration of conformers lacking the secondary structure criteria needed for the pseudoknot nucleation, which can be for e.g. the presence of the P1 helix, etc. This class may involve unfolded RNA or conformers having a secondary structure that prevents the binding of both the site-specific Mg^2+^ and SAM. This population or species of conformers are considered as *binding incompetent*.

**Figure 2.**
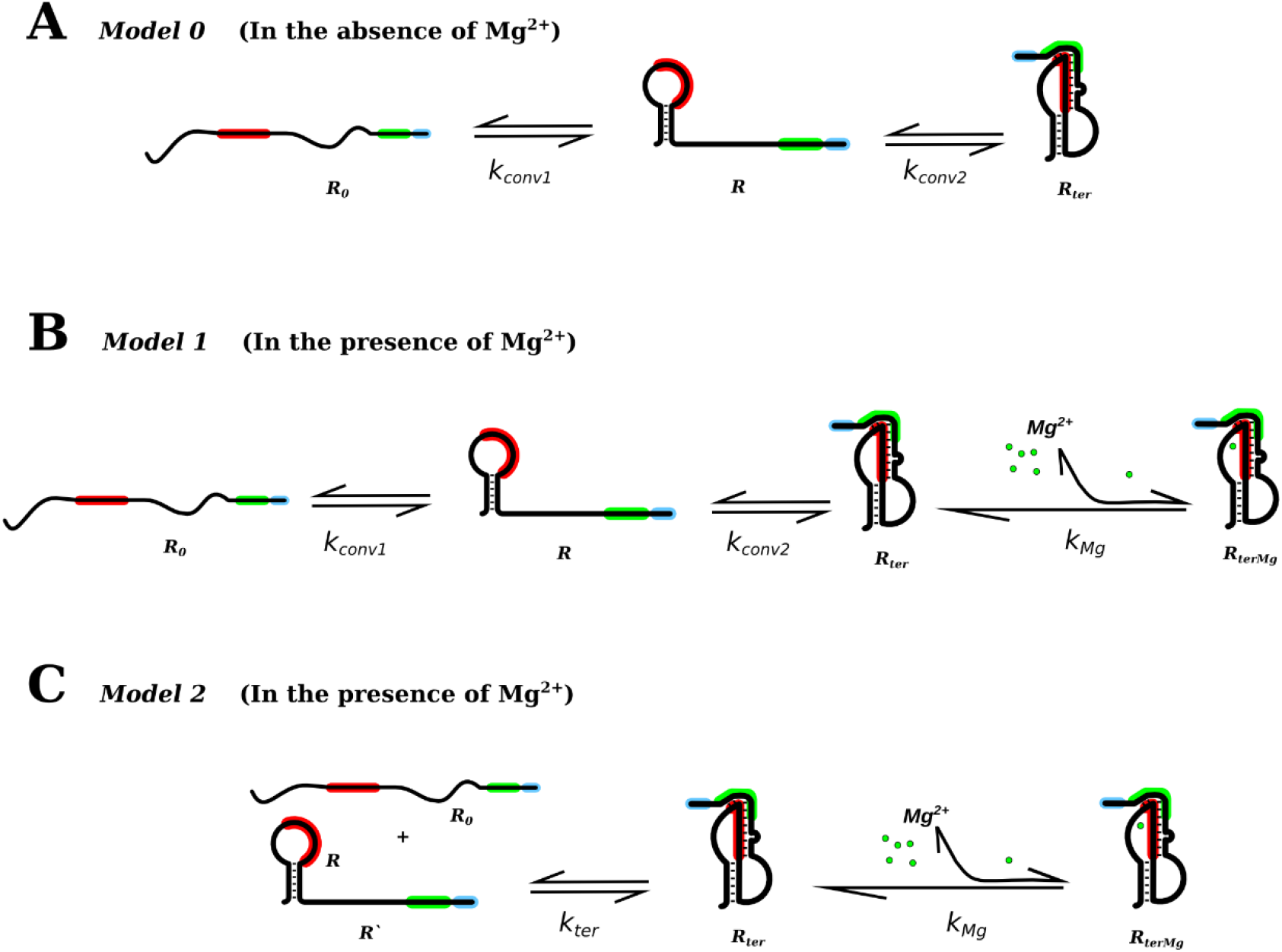
Models for Mg^2+^ binding and pseudoknot nucleation resulting in SD sequence sequestration. The figure illustrates equilibrium models for the RNA folding and Mg^2+^ binding (**A**) in the absence of Mg^2+^ (designated as *Model 0*) and (**B**) in the presence of Mg^2+^ (designated as *Model 1*). The schematic diagram in (**C**) shows a reduced model (designated as *Model 2*) that is convenient to use for practical predictions and direct comparison with experiments. The symbols for the RNA species/population and the equilibrium association constants are illustrated below their corresponding RNA skeletons and reaction arrows, respectively.

*R* is the concentration of the riboswitch RNA having the specific secondary structure criteria needed for the pseudoknot nucleation, which could be for e.g. conformers forming the P1 helix, L1 loop, etc. This RNA population can further fold to form the pseudoknot. Conformers belong to this class are also considered to be *binding incompetent*. The secondary structure criteria mentioned here are general examples. More details regarding the nature of the specific secondary structure criteria will be discussed later in this manuscript.

*R*_*ter*_ is the concentration of riboswitch having the specific secondary structure criteria needed for the pseudoknot nucleation and possesses a nucleated or partially formed pseudoknot. Hence, the latter conformer population is considered as *binding competent*.

*R*_*terMg*_ is the concentration of Riboswitch that resembled *R*_*ter*_ after binding with Mg^2+^ ions. The latter conformer population is considered as *binding competent* and constitutes the *fraction bound*.

*Mg* is the concentration of free magnesium ions.

To be able to calculate the fractions of the various species, the following quantities are defined, *N* specifies the stoichiometry. In other words, it is the number of moles of Mg^2+^ ions that specifically binds with the 1 mole of the RNA, to stabilize of the nucleation of the pseudoknot. Because the model describes site-specific binding, *N* is a value that refer to the number of ions that is bound to a single RNA molecule. Indeed, atoms are countable, and therefore in this model, the minimal value for *N* is unity and is only physically sensible if restricted to integer values. For this riboswitch, no distinct multiple transitions were observed upon the addition of Mg^2+^ ions [19], as had been the case for other riboswitches (e.g. the SAM-I riboswitch [5]). That is, only a single additional state was observed in the presence of magnesium compared to that were seen in the absence of these ions. Hence the possibility of sequential or independent mechanisms [23] of multiple ion-binding was excluded. Therefore, the cases of *N* > 1 considered here refer to the simultaneous binding of multiple ions. Though the latter is certainly not a common scenario yet remains a possibility to explore.

*Mg*_*tot*_ and *R*_*tot*_ are designated to the total Mg^2+^ ion and total RNA concentrations, respectively.

At equilibrium, and assuming a *N*: 1 magnesium to RNA stoichiometry, the following equilibrium constants can be defined,

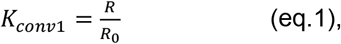

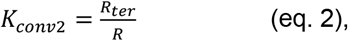

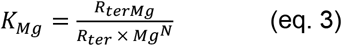

*where,*

*K*_*conv*1_ is the association constant for the conversion of *R*_0_ to *R* in the absence of Mg^2+^, *K*_*conv*2_ is the association constant for the conversion of *R* to *R*_*ter*_, also in the absence of Mg^2+^, and *K*_*Mg*_ is the binding association constant of Mg^2+^ to *R*_*ter*_ and hence forming *R*_*terMg*_.

To reduce the number of initial parameters needed to apply the model, and to be able to directly use the model to compare with experimental data without making any assumption about the secondary structure, *Model 1* is reduced to *Model 2* (**Figure 2C**), *where*,

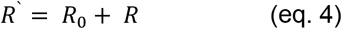

We therefore can define the equilibrium constant, *K*_*ter*_, as the apparent association constant for the conversion from *R*′ to *R*_*ter*_ in the absence of Mg^2+^, which is given by,

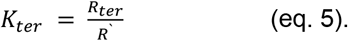

Structurally, *K*_*ter*_ in **eq. 5** represents the association constant for the nucleation of the pseudoknot. Whereas, *K*_*Mg*_ is the equilibrium constant for the binding of Mg^2+^. The quantity *R*_*ter*_ + *R*_*terMg*_, which represents the concentration of RNA having a partially or fully formed pseudoknot (bound or unbound to Mg^2+^), had been determined experimentally, at several Mg^2+^ concentrations [3].

By substitution for *R*_*ter*_ and for *Mg*^*N*^in (**eq. 3**) it can be shown (**Supplementary methods**), for *N* = 1, that the concentration of the RNA bound to Mg^2+^ (*R*_*terMg*_) can be calculated from a quadratic, for which the solution is given by the expression,

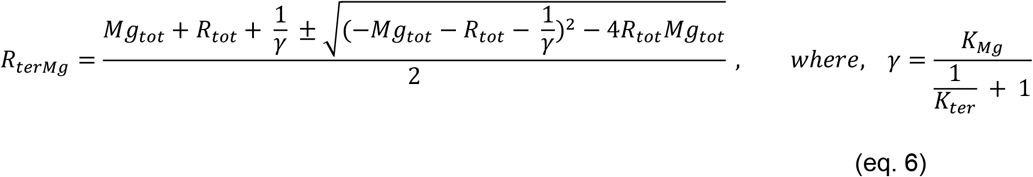

The reader may refer to the **Supplementary methods** for the details of the derivation. Similarly, for *N* = 2, *R*_*terMg*_ can be calculated as a function of *Mg*_*tot*_ and *R*_*tot*_ from a cubic equation which can be solved numerically (see **Supplementary methods** for the details of the derivation).

### 2.2 Determination of equilibrium constants from experimental data

In the presented model, the two association constants (*K*_*ter*_ and *K*_*Mg*_) in (**eq. 6**), were determined from experimentally reported values in the literature [3]. In these NMR Chemical Exchange Saturation Transfer (CEST) experiments, the base pairing interaction of G22 and C43 was monitored, and the fraction of the RNA having this base pair was reported in the absence and presence of different concentrations of Mg^2+^ ions. These measurements therefore can provide solid bases for determination of equilibrium constants. For clarity, it is worth noting that throughout the manuscript, all discussed free energy terms refer to the *standard* free energies.

The two constants were initially determined from single experimental points (**Table S1-S3**) for both *N* = 1 and *N* = 2, to examine the calculated standard free energy range obtained at different data points. The equilibrium constants were then determined by fitting to all data points i.e. using a global fit. Global fitting was done by varying the standard free energies and the corresponding association equilibrium constants (*K*_*ter*_ and *K*_*Mg*_) were calculated from 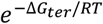 and 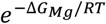, respectively. The latter constants are then used to calculate the probability of the riboswitch RNA having the nucleated pseudoknot *R*_*ter*_ + *R*_*terMg*_ using a total RNA *R*_*tot*_ similar to what had been used in the experiment (0.3 mM) and using total magnesium concentrations identical to those used in the NMR experiment which are 0, 0.05, 0.25, 0.5 and 2.0 mM. The best fits were determined by values of free energies that minimized *Chi* squared (*Chi*^2^), which is obtained from the difference between calculated and experimental values and their corresponding experimental errors, as given by,

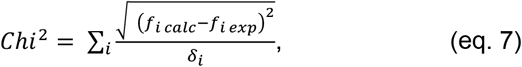

where, *f*_*i calc*_ and *f*_*i exp*_ are the calculated and experimental fractions (probability), respectively, of the riboswitch RNA having the nucleated pseudoknot for data point *i*. The sum here is taken over all experimentally determined points and takes the experimental errors *δ*_*i*_ into account. It should be noted that the two constants were fitted to the *Model 0* (to calculate *K*_*ter*_) and *Model 2* (to calculate *K*_*Mg*_ and refine for *K*_*ter*_) simultaneously.

As detailed in the next section, once these two constants have been determined by fitting, the final concentrations of all RNA species of the model can be computed for any given total concentration of magnesium, e.g. to model titration curves, using equations (**6** and **3** & **5**). The apparent standard free energy change of the system for each of the two assumed stoichiometry, can be therefore estimated from the sum of the two standard free energy terms, that is,

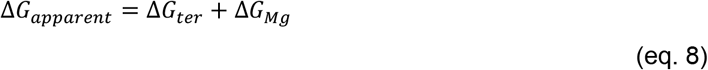

The non-specific binding of Mg^2+^, is assumed to equally contribute to all the RNA species considered in the model, and hence not need to be accounted for when fitting the binding constant. Hence for simplicity, it was neglected initially when obtaining the model parameters. However, the experimentally observed apparent free energy change of the system is given by,

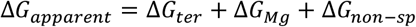

Where, Δ*G*_*non*−*sp*_ is the binding free energy contribution of Mg^2+^ ions that are non-specifically bound to the RNA surface.

By definition of equilibrium (**eq. 3**), the apparent dissociation constant which is given by, 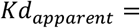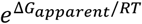, is based on the *free* magnesium concentration. Often however, the experimentally determined value for the apparent dissociation constant, is reported in the literature in terms of the [*Mg*_*tot*_½], which is half of the *total* magnesium concentration at which the change occurs. Both quantities can be readily calculated from the model.

### 2.3 Estimating the fraction forming the pseudoknot inside the cell

The level of magnesium inside the cell cytoplasm is assumed to be constant, since it is likely to be actively maintained by Mg^2+^ transporters ([24] and references therein). Hence, it can be assumed that the concentrations of the *free* magnesium and the *total* magnesium are equal when considering an in-cell model i.e. *Mg* ≅ *Mg*_*tot*_ ≅ *Mg*_*cyt*_. The latter quantities would in this case refer to the bulk accessible (i.e. free) and maintained Mg^2+^ concentration level in the cytoplasm, and the symbols *Mg*_*tot*_ and *Mg*_*cyt*_ are used interchangeably for the in-cell model throughout the manuscript.

Thus, for *N* = 1, *R*_*terMg*_ is given by, (see **Supporting methods** for stepwise derivation),

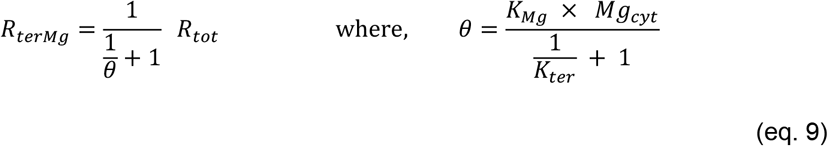

This simplifies calculating the fraction of *R*_*terMg*_, 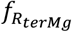, which can be obtained directly from the equilibrium constants and the bulk magnesium concentration, and it is given by,

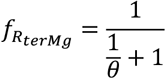

### 2.4 The incorporation of RNA folding energetics into the model and simulating titraton curves of mutants

To be able to incorporate RNA folding energetics into the model, *Model 1* needs to be applied (as illustrated in **Figure 2B**). However, an assumption must be made about the minimal criteria of the secondary structure required to enable the pseudoknot to be formed. Consequently, the equilibrium constant for RNA secondary structure formation (*K*_*conv*1_), can be determined, and from which *K*_*conv*2_ is computed (**eq. 1** and **2**). Specifically, in the absence of magnesium ions, it can be shown that,

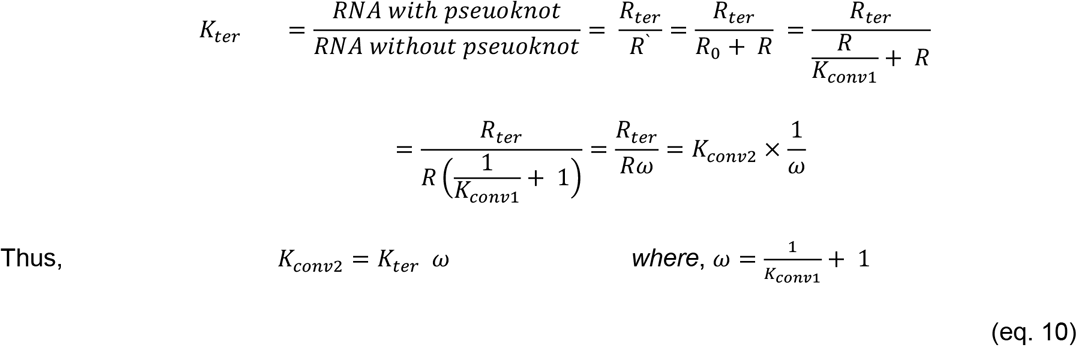

**Eq. 10** shows that *K*_*conv*2_ can be calculated from the experimentally determined *K*_*ter*_ and from the RNA energetics (which determine *K*_*conv*1_), if an assumption about the secondary structure is made. For instance, *K*_*conv*1_ can be calculated from the energies of individual conformers containing a specific substructure, and Δ*G*_*conv*1_ is readily computed from,

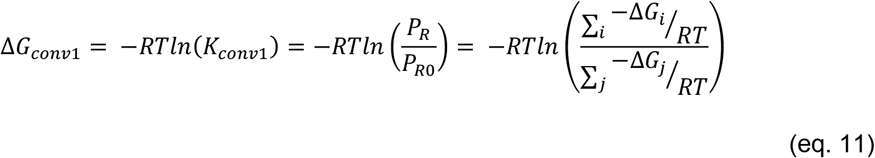

where, the probabilities *P*_*R*_ and *P*_*R*0_, are the sum of the probabilities of all suboptimal conformers in which a specific substructure is present (denoted as conformers *i*) or absent (denoted as conformers *j*), respectively. *R* and *T* are the gas constant and temperature in Kelvin, respectively.

The equilibrium constant here is calculated in the absence of any tertiary interactions, and in the absence of any ligands. In this work, we have used the computational method and code previously reported in reference [25], using optimal (i.e. minimal free energy) and all suboptimal conformers (obtained from the *ViennaRNA* Package, using the *RNAsubopt* program [26]) with an energy range up to 11 kcal/mol higher than the minimal free energy conformer and a temperature of 298.15 Kelvin.

Titration curves for mutants were calculated by determining their corresponding *K*_*conv*1_, for the fixed and experimentally determined constant, *K*_*conv*2_ and *K*_*Mg*_, assuming specific secondary structure criteria for conformers that are pseudoknot competent. The presence of unpaired motif sequences that may form a nucleus for the triplex/pseudoknot formation was taken as the condition for the secondary structures that will be involved in the subsequent step. Namely, secondary structures with unpaired nucleotides (C23 and A24) and their interacting partner nucleotides (U40, A41 and G42) were taken to be the population that constitute the RNA species *R*, which is thought to be able to proceed to form a pseudoknot. *Model 1* was then applied to predict the fraction of RNA that may have a nucleated (partially formed) pseudoknot by using **eq. 10** to determine the mutant/sequence specific *K*_*ter*_. In these mutants, *K*_*conv*2_ and *K*_*Mg*_ are considered fixed because there was no change in the nucleotides that are directly involved in the pseudoknot formation nor those that are in direct contact with SAM in the crystal structure.

## 3 RESULTS

### 3.1 The SAM-II riboswitch exhibits a favorable Mg^2+^ binding energy and an unfavorable pseudoknot nucleation free energy

As described in the methods section, for a given *N*, the two equilibrium constants (*K*_*ter*_ and *K*_*Mg*_) used in **eq. 6** for determination of the fraction of RNA having a nucleated pseudoknot, can be computed from a single measurement in the absence of Mg^2+^ along with each data point independently (i.e. single point data or single measurement in the presence of Mg^2+^) or can be fitted to all data points simultaneously (global fit).

For instance, at a Mg^2+^ concentration of 2.0 mM, the equilibrium constants and concentrations of all RNA species can be calculated from a single experiment reporting the fraction of RNA that possesses a pseudoknot like structure in the presence and absence of Mg^2+^ (i.e. from a single point data) as follows. The probability of riboswitch RNA having the nucleated pseudoknot, in the absence of Mg^2+^ (i.e. the probability of the species *R*_*ter*_) was determined experimentally, using NMR [3] by monitoring the C43-G22 base pairing interaction, and was found to be 0.093 (9.3%). The latter experiments were done using 0.3 mM RNA (*R*_*tot*_). Since in the absence of Mg^2+^ (i.e. in *Model 0*) the total RNA is given by, *R*_*tot*_ = *R*′ + *R*_*ter*_, hence the probability of *R*′ and the value of *K*_*ter*_ (from **eq. 5**) are 90.7% and ~0.103, respectively. This corresponds to a free energy of ~ 1.35 kcal/mol. In the presence of magnesium (i.e. in *Model 2*), at a concentration of 2.0 mM, the probability of the riboswitch RNA having the nucleated pseudoknot (data were obtained by monitoring the same base pair), representing the probability of the species *R*_*ter*_ + *R*_*terMg*_, was reported in the same NMR experiment [3] to be 22%. This implies that the probability of *R*′ is equal to ~78%.

Given that the total RNA in the presence of magnesium ions is equal to,

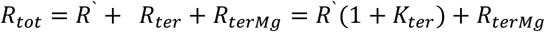

and that *K*_*ter*_ is known, then, the concentration and probability of *R*_*terMg*_ are ~0.04 mM and ~13.8%, respectively. Further, since *Mg*_*tot*_ = *Mg* + *N*. *R*_*terMg*_, the amount of free magnesium for *N* = 1, *Mg*, is obtained from *Mg* = *Mg*_*tot*_ − *N*. *R*_*terMg*_ = 1.96 mM. The magnesium binding constant, *K*_*Mg*_, is therefore determined from **eq. 3**. The latter was found to be *K*_*Mg*_ = 877.5 M^−1^. The obtained values for the same equilibrium constants (*K*_*ter*_ and *K*_*Mg*_) by global fitting are shown in **Table 1** and **Table S4** for *N* = 1 and *N* = 2, respectively.

**Table 1.** Model parameters obtained by global fitting for 1:1 stoichiometry (*N* = 1). The table on the left shows the model parameters, i.e. equilibrium constants and corresponding free energies, that were obtained from global fitting to experimental data (*Model 2*), with *N* = 1. The table on the right lists parameters used in *Model 1*, taking the presence of unpaired C23, A24, U40, A41 and G42 as the criteria for the nucleation of the pseudoknot (see **Methods** section for further details). The latter condition in the secondary structures means that contacts involved in the pseudoknot nucleation are accessible, allowing the interaction (i.e. base pairing) between CA (nucleotides 23:24) and UAG (nucleotides 40:42) to occur.

| Model<br>Parameter<br>(Model 2) | Values<br>($N = 1$)<br>*in vitro* | Values<br>($N = 1$)<br>*In-cell* | Model<br>Parameter<br>(Model 1) | Values<br>($N = 1$)<br>*in vitro* | Values<br>($N = 1$)<br>*In-cell* |
| --- | --- | --- | --- | --- | --- |
| $K_{ter}$ | 0.103 | 0.103 | $K_{conv1}$ | 22.54 | 22.54 |
| $Kd_{ter}$ | 9.75 | 9.75 | $\Delta G_{conv1}$ | -1.85 kcal/mol | -1.85 kcal/mol |
| $\Delta G_{ter}$ | 1.35 kcal/mol | 1.35 kcal/mol | $K_{conv2}$ | 0.107 | 0.107 |
| $K_{Mg}$ | 878 M<sup>-1</sup> | 859 M<sup>-1</sup> | $\Delta G_{conv2}$ | 1.32 kcal/mol | 1.32 kcal/mol |
| $Kd_{Mg}$ | 1.14 mM | 1.16 mM | | | |
| $\Delta G_{Mg}$ | -4.02 kcal/mol | -4.00 kcal/mol | | | |
| $[Mg_{tot}^{1/2}]$ | 10.0 mM | 10.2 mM | | | |
| $\Delta G_{apparent}$ | -2.61 kcal/mol | -2.61 kcal/mol | | | |
| $\chi^2$ | $15.65 \times 10^{-3}$ | $15.70 \times 10^{-3}$ | | | |
| $\chi^2_{err}$ | 1.475 | 1.480 | | | |

Both free energies obtained from single points data and from global fitting to all data points show similar trends. Indeed, the free energy of the pseudoknot (partial) formation is unfavorable whereas the stability of the pseudoknot comes from the contribution of the Mg^2+^ ion, which binds with a free energy of ~ − 4.0 kcal/mol, assuming a 1:1 stoichiometry.

### 3.2 Magnesium ions binds to a specific site within the SAM-II Riboswitch with a 1:1 stoichiometry

To be able to confirm the stoichiometry of the RNA to Mg^2+^ binding, the model was derived with the two assumptions, *N* = 1 and *N* = 2 (see **Supplementary methods** for the detailed derivation and solution). The free energies of the model were calculated from experimental data (from either single points or global fit). In the previously reported SEC experiment [7], in which the reduction of the size (i.e. compaction) of the riboswitch was monitored, showed that half of the total magnesium concentration [*Mg*_*tot*_½] at which this change occurs is ~ 6 mM. The latter corresponds to an apparent free energy change (Δ*G*_*apparent*_) of ~ − 3.0 kcal/mol, if a 1:1 binding is assumed. Since it is likely that a part of the magnesium ions would be non-specifically bound with a favourable free energy contribution, then the apparent dissociation constant *Kd*_*apparent*_ (which is estimated experimentally by [*Mg*_*tot*_½]) of the site-specific bound magnesium should be ≥ 6 mM, and unlikely to be less. The latter value could be thought of as a condition for determining the correct stoichiometry.

The global fitting of the model parameters based on both *N* = 1 and *N* = 2, revealed in a similar pseudoknot formation association constant *K*_*ter*_ of 0.103 and 0.105, respectively, (**Tables 1 and S4**, respectively), which correspond to free energies, Δ*G*_*ter*_ of 1.35 and 1.34 kcal/mol, respectively. Thus, with both assumptions the pseudoknot formation is not an energetically favourable process. The binding association constant on the other hand, of RNA to Mg^2+^, with 1:1 stoichiometry (i.e. *N* = 1) and 1:2 stoichiometry (i.e. *N* = 2) were found to be 8.78 × 10^2^ M^−1^ and 4.51 × 10^5^ M^−1^, respectively. These values correspond to binding free energy of −4.0 kcal/mol and −7.1 kcal/mol, respectively. This makes the apparent free energy change, Δ*G*_*apparent*_, (**eq. 8**) for the 1:1 stoichiometry model is −2.61 kcal/mol, corresponding to a [*Mg*_*tot*_½] = 10 mM, while for the 1:2 stoichiometry is −6.32 kcal/mol, corresponding to a [*Mg*_*tot*_½] = 4.54 mM. It is therefore clear that the SAM-II riboswitch is likely to bind with magnesium ions with 1:1 stoichiometry (*N* = 1) since its [*Mg*_*tot*_½] is > 6 mM, whereas *N* ≥ 2 do not fulfil the condition for correct stoichiometry mentioned earlier.

Further, in all the attempts to fit the data to *N* = 2 (**Figure S2A**), both single point determination of the model parameters and global fits revealed in a *Chi*^2^ that is higher than it is for *N* = 1 (**Figure 3A**) and would not be considered as a good fit (**Tables 1** and **S4**). For instance, the model in this case predicts that the fraction of RNA having a pseudoknot, at 0.25 mM Mg^2+^, is only ~ 9.7 %, which is lower than in the case of *N* = 1 ( which predicts a fraction of ~ 11.1 %) and lower than the reported experimental value (~ 11 %) [3]. This implies that if the RNA specifically binds with two Mg^2+^ ions, we would not see any significant increase in the probability of the pseudoknot in the presence of 0.25 mM magnesium. This is clearly inconsistent with the reported experimental value and conclusion [3]. The *R*_*terMg*_ titration curve in the case of *N* = 2 is found to shift to the right and become steeper compared to the case of *N* = 1 (**Figure S2B** and **3B**, respectively). These are obvious changes for higher binding stoichiometry. The shift is due to the naturally low stability of the pseudoknot in the absence of Mg^2+^, and with stoichiometry *N* = 2, the minimal number of Mg^2+^ ions needed to saturate the RNA had now increased, which means that higher magnesium concentration is needed to induce binding. On the other hand, the increase in steepness is due to the lower binding energy when calculated with 1:2 RNA to Mg^2+^ resulting in that the bound RNA becomes stable and dominate the population as soon as Mg^2+^ ions concentration reaches a certain lower limit. The same applies to the titration curve for the fraction of binding competent conformers (i.e. *R*_*ter*_ + *R*_*terMg*_).

**Figure 3.**
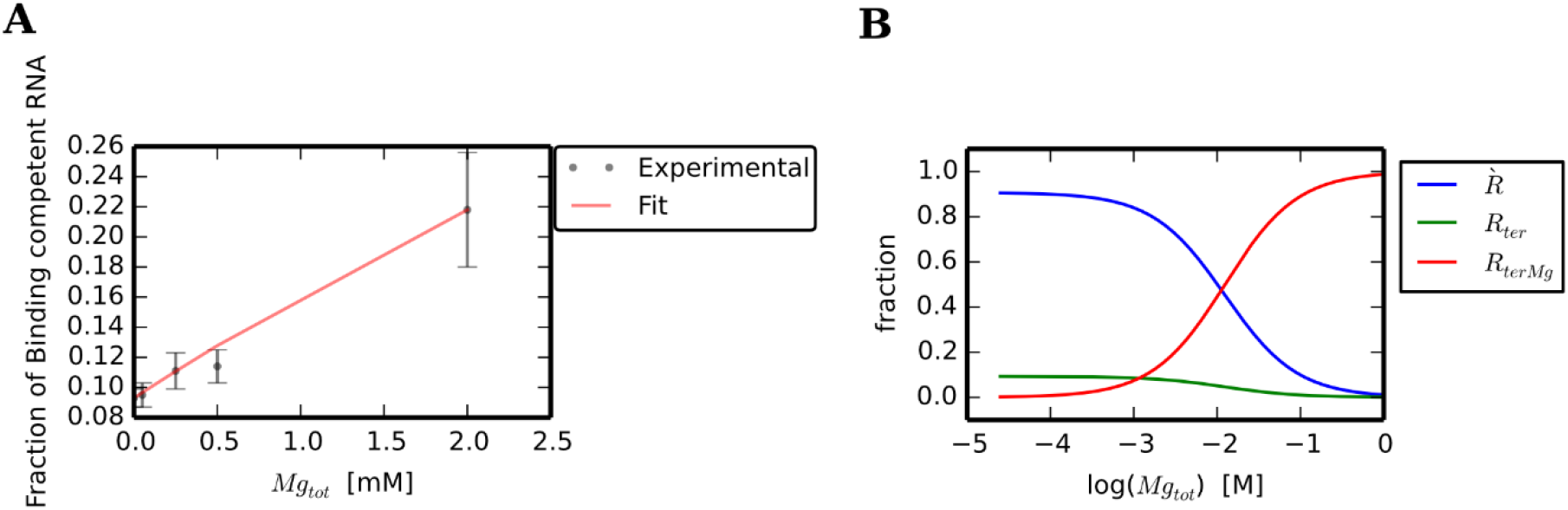
Fractions of various RNA species as a function of magnesium concentration assuming a 1:1 RNA to Mg^2+^ stoichiometry. Panel (**A**) shows a plot for the experimental data (and their associated error bars) versus the calculated fraction of binding competent conformers, using *N* = 1. The calculated fraction is obtained by the global fitting of model parameters to the fraction of RNA with a nucleated pseudoknot inferred from NMR CEST experimental data and reported by Chen et al [3], in which the base paring of G22 and C43 was monitored. Panel (**B**) shows a plot of the calculated fraction of each RNA population/species at an increasing magnesium concentration calculated from the globally fitted parameters for *N* = 1. The fractions of each of *R*′, *R*_*ter*_ and *R*_*terMg*_ (fraction bound) as a function of magnesium concentration (on a logarithmic scale) are shown in blue, green and red, respectively. From structural point of view, *R*′ and fraction binding competent (*R*_*ter*_ + *R*_*terMg*_) represent the fractions of RNA without and with a nucleated pseudoknot, respectively.

Indeed, based on the latter calculations for *R*_*terMg*_, it can be inferred that increasing *N* to > 2 would lead to the increase of the steepness of the titration curve, and the curve will shift further to right, in a similar manner as seen when *N* was increased from 1 to 2. This would mean that the Mg^2+^ concentration will need to be even higher (≫ 0.25) to induce any noticeable changes in the RNA conformation. The latter expectations can be seen from calculations for *N* = 3,4, 5 … etc., using the in-cell model (data not shown). Cleary, this is unlikely to explain the reported experimental observations. These arguments verify that, the stability of the nucleated pseudoknot formed by the SAM-II riboswitch is increased by specifically binding to Mg^2+^ ions with 1:1 ratio and it is unlikely that the increase in the latter number of Mg^2+^ ions to the RNA would improve the agreement between the experimentally determined and calculated fractions, based on site-specific binding.

### 3.3 Prediction of the fractions forming a pseudoknot confirms the plausibility of the model

Using the SAM-II riboswitch as a case study, a simple equilibrium binding and mechanistic model was constructed, where its equilibrium constants were determined based on experimental observations from the literature. Despite its simplicity, the model predicted experimental observables based on equilibrium constants that can be, in principle, determined from even a single data point and consequently be used in the prediction of titration curves for the pseudoknot formation.

Specifically, it was proposed that magnesium ions stabilize the nucleated pseudoknot with a binding stoichiometry of RNA:Mg^2+^ of 1:1. The model constants (*K*_*Mg*_ *and K*_*ter*_), and their corresponding free energies were derived from experimental observations reported in the literature (see methods for details). Namely, the fraction of RNA having the pseudoknot (i) in the absence of magnesium (**Table S1**) and (ii) in the presence of a known total concentration of magnesium. The concentrations of free and bound Mg^2+^ were calculated (as described in the **Methods** and **Supplementary methods**). A summary of final parameters (i.e. globally fitted) and concentrations of all RNA and Mg^2+^ ions species are listed in (**Table 1 and Table S1**).

The pseudoknot nucleation/formation association constant *K*_*ter*_ was found to be 0.103 (**Table 1**), which corresponds to a free energy of 1.35 kcal/mol. Thus, the pseudoknot formation is not an energetically favourable process. The binding association constant on the other hand, for Mg^2+^ ions to the RNA with 1:1 stoichiometry (i.e. *N* = 1), was found to be 8.78 × 10^2^ M^−1^ which corresponds to a binding free energy of −4.02 kcal/mol.

The obtained values of the equilibrium constants were used to calculate the concentration and fraction of the RNA bound to Mg^2+^ from the derived expression in (**eq. 6**). The expression can be used to predict titration curves using the proposed two step-model. The concentration of the magnesium ions bound to the riboswitch *R*_*terMg*_ and its corresponding fraction were calculated as a function of magnesium concentration, given the total RNA concentration. Consequently, all other RNA species (**Figure 3B**) were determined as a function of magnesium concentration (from equations **1**–**3** and **5**). **Figure 3A** shows a reasonable agreement between the calculated and the experimentally determined fractions of the RNA with a fully or partially formed pseudoknot, at equilibrium.

### 3.4 The impact of magnesium ions concentrations on the SAM-II riboswitch inside the cell

Since Mg^2+^ ions level is regulated and maintained constant by other means in the cell [27–29], it could be assumed that the decrease in the concentration level of the free Mg^2+^ due to the consumption during binding with other molecules including RNA, is re-compensated. On this basis, it can be shown that, the fraction of RNA that would bind to magnesium is dependent on: a fixed Mg^2+^ concentration, the stoichiometry, and on the equilibrium constants (*K*_*Mg*_ *and K*_*ter*_) and is independent of the total RNA concentration (see methods **eq. 9**). Hence, this approximation effectively means that, at a given Mg^2+^ in the cell, the fraction of riboswitch RNA that is SAM binding competent in a cell will remain fixed despite the total concentration of the riboswitch RNA.

Several studies reported that the total magnesium in bacterial cells such as *E. Coli* ranges between 20 [30] and 100 mM [31], whereas the concentration of the free magnesium in the cytosol is thought to be much lower, and estimated to be only within the range of 1-2 mM [32,33]. The fraction of the riboswitch that is binding competent (i.e. the probability of conformers with a pseudoknot which is *R*_*ter*_ + *R*_*terMg*_), based on **eq. 9**, is therefore estimated to be between ~ 16.0% and ~ 21.8%.

In general, the calculated *fraction of binding competent conformers* as a function of Mg^2+^ concentration with a 1:1 binding (*N* = 1), is very similar for the in vitro and in-cell models (**Figures 4**). A similar observation is seen for *N* = 2, with only a slight difference is noticeable between the in vitro and in-cell models. The same applies for the calculated *fraction bound* (**Figure S3**). In other words, the predicted titration curves for the in vitro and in-cell models are very similar, illustrating that the assumption about the in-cell concentration of magnesium is plausible and can be used as a convenient approximation to study riboswitch folding for both in-cell and in vitro, provided that in the latter case, the condition *Mg*_*tot*_ ≫ *R*_*tot*_ holds. However, **eq. 9** shows that at fixed cellular concentration of Mg^2+^ ions, the concentration of *R*_*terMg*_ varies linearly with *R*_*tot*_, which is significantly different than the expected behavior in vitro, where the concentration of the bound complex *R*_*terMg*_ would be limited by the maximum concentration of magnesium (**Figure S4**).

**Figure 4.**
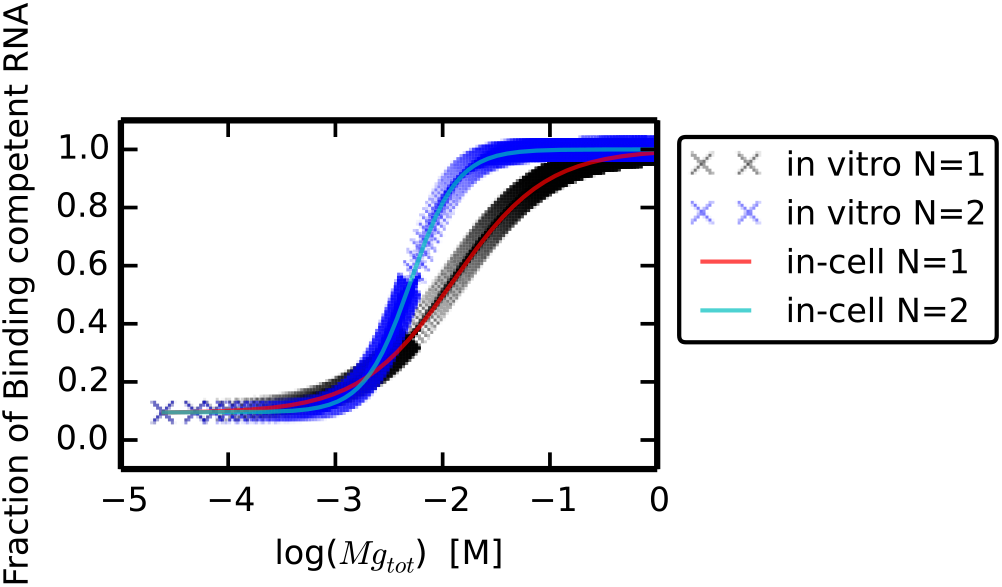
The *fraction of binding competent conformers* calculated with various model assumptions. The figure shows the *fraction of binding competent conformers* i.e. (*R*_*ter*_ + *R*_*terMg*_), plotted as a function of the total magnesium molar concentration for *N* = 1 and *N* = 2, each for both in vitro and in-cell approximations. The logarithmic plot enables the comparison between the probability of the pseudoknot as predicted by the various assumptions.

To illustrate the effect of **Eq. 9**, titration curves were calculated for a fixed *R*_*tot*_ with a varying *Mg*_*tot*_ and vice versa for both in vitro and in-cell models. In panels (**A**) and (**B**) of **Figure S4**, the concentrations (and fractions) were calculated at increasing total Mg^2+^ concentrations using the in vitro model, at different total RNA concentrations (which are, for both plots, denoted by the ligand on the right). The calculated fractions in (**A**) are similar (overlapping) for the different total RNA concentrations. Very similar results (almost visually indistinguishable) were obtained for the corresponding titration curves when the in-cell model was used (data not shown). In each of the curves shown in the latter plots, *R*_*tot*_ was constant.

Titration curves in panels (**C**) and (**D**) of the same figure (**Figure S4**), show the calculated fraction of binding competent RNA and bound RNA concentration, respectively, using a limited concentration of a 1 mM Mg^2+^, for an increasing total RNA concentration. Unlike the predicted curves using the in vitro model (depicted in red), in which the bound RNA concentration and fraction are limited by the Mg^2+^ ions (i.e. reaching a plateau), the in-cell model however shows that the bound RNA increases linearly with the increase of the total RNA. Hence, the fraction of bound and binding competent RNA will remain fixed in the cell, despite the amount of expressed RNA. This is mainly because in the cell the consumed Mg^2+^ ions are recompensated. Whereas in vitro, the increase of the bound RNA will consume the Mg^2+^ ions eventually limiting the increase of the concentration of the bound complex (panel **D**), and thus leading to the apparent decrease in the fraction of bound and binding competent RNA, as shown in (panel **C**). In other words, in the cell where Mg^2+^ ions hemostasis is maintained, the fraction of RNA having a pseudoknot is constant, regardless of the total RNA expressed. While in vitro, as the total RNA concentration increases, the fraction of the unbound RNA species will increase, which is expected given the limited concentration of Mg^2+^ ions.

### 3.5 Linking secondary structure energetics to ligand bindng and examining the plausible secondary structure conditions that define the species *R*

Determination of the model association constants *K*_*ter*_ and *K*_*Mg*_ involved in *Model 2* makes it possible to then use *Model 1* (**Figure 2**) to gain further insights into the system and study the behavior of different mutants. To accomplish such task, *K*_*conv*1_ and *K*_*conv*2_ need to be determined. The computation of the equilibrium constant of the conversion of the secondary structures to tertiary (which was designated as *K*_*conv*2_) can be determined based on an assumption about the minimal criteria of the secondary structure which is needed to be in a conformer to be able to form a pseudoknot. Once plausible criteria are chosen, the secondary structure energies of those conformers can be used to calculate *K*_*conv*1_ (from **eq. 11**), and hence *K*_*conv*2_ can be estimated (from **eq. 10**) for the experimental sequence (showed in **Figure 1**). The latter sequence was used in the NMR study reported in [3], which had two point mutations with respect to the wild sequence.

In order to check the plausibility of the different secondary assumptions, the apparent free energy change for different minimal secondary structure criteria needed for the pseudoknot formation was examined for the crystallized sequence. It had been shown that this sequence binds to SAM with a higher free energy compared to the experimental sequence, showing a discrepancy of ~ 0.4 kcal/mol [14]. Although SAM binding was not modelled in this study, plausible secondary structure criteria should provide a similar trend, i.e. the apparent free energy of binding should increase with the crystal sequence compared to that of the experimental sequence. **Figure S5** illustrates various criteria that could be examined, including a nucleated pseudoknot and a nucleated P1 helix.

To assess the plausibility of such assumptions, the secondary structure conditions illustrated in **Figure S5** and listed in **Table S5**, that essentially define the population *R* in *Model 1,* were used to calculate the magnesium titration curves by monitoring the fraction forming a pseudoknot, for both the experimental and crystal sequences using *Model 1*. A secondary structure condition was considered plausible, if the crystal structure sequence shows an apparent free energy change (i.e. an apparent Mg^2+^ binding or equivalently midpoint transition [*Mg*_*tot*_½]) that is higher compared to the unmutated experimental sequence (i.e. lower apparent binding), and hence would be consistent with the fact that the crystal structure sequence showed a lower binding affinity to SAM using Isothermal calorimetry (ITC) [14].

Although the actual apparent binding with SAM is not calculated by this model, the apparent binding with Mg^2+^, or similarly the equilibrium constant *K*_*conv*1_, is taken as an indicator for the efficiency of binding with SAM. This simple check allowed to test whether an assumption about the secondary structure is consistent with the experimental finding. Although cannot confirm a specific condition, the test can at least exclude conditions that would give contradictory results with the ITC experimental measurement, and hence providing new insights into the folding mechanism. These conditions can be thought of as plausible reaction pathways through which the reaction may proceed.

For instance, as shown by the plot in **Figure S6A**, the *L3_nucleated_pseudoknot* condition (illustrated in **Figure S5**), is not a valid secondary structure assumption, because of the increased stability of the P1 helix of its crystal sequence, as indicated by the high *K*_*conv*1_ and hence the low apparent Mg^2+^ binding energy, compared to the experimental sequence. This implies, that if this condition was possible, then the crystal sequence variant would have shown an apparent binding affinity with SAM that is higher than the experimental, a prediction that clearly contradicts with the experimental finding. Thus, this secondary structure condition can be excluded. The plot in **Figure S6A** illustrates that several criteria that could provide similar trends, including a nucleated pseudoknot and a nucleated P1 helix. Though qualitative, and does not verify a specific folding pathway, the analysis has proven to be useful. Further experimental data, and the incorporation of SAM binding, would be needed to verify (and perhaps quantify) the dominating folding pathway(s) in which the molecule would go through from its secondary to the tertiary structure. For the next part of the manuscript, a nucleated pseudoknot (as specified in the methods section and described in **Table S5** and **Figure S5**), was chosen to be used for further calculations.

### 3.6 The stability of the P1 helix in the wild sequence correlates with the fraction of binding competent conformers

The resulting parameters, based on the presence of nucleated pseudoknot, for the experimental sequence are shown in **Table1.** The constant *K*_*conv*2_ can be then fixed in calculations of titration curves for mutants or sequences with changes that are not involved in the pseudoknot nor in the site-specific Mg^2+^ binding. These mutants or sequences include for example, the unmutated wild sequence, crystal structure sequence (which incorporates the mutations of two base pairs in the P1 helix), longer sequences to incorporate the start codon and various mutations in the P1 helix, among many others. As an example, calculated magnesium titration curves for the wild sequence (length=52 nucleotides), experimental, crystal structure and the 56 nucleotide-long wild sequence are illustrated in (**Figure S6B**). The study of the energetics and ensemble populations of such sequences may offer new insights on the structural mechanism and function of the riboswitch.

**Figure 5** shows the concentrations of various RNA populations used in *Model 1* for the wild sequence as a function of the total magnesium concentration (*Mg*_*tot*_). The titrations curves were computed with a total RNA concentration of 0.3 mM for transcript lengths 52 and 56 nucleotides (panels **A** and **B**), respectively. It is evident that, as the sequence becomes longer, the concentration of *R* becomes smaller due to the competition with other possible conformers, as opposed to the concentration of *R*_0_ which increases with the longer RNA sequence, indicating the increase of the conformers that are incompetent for pseudoknot formation. These findings are generally consistent with the fact that the number of possible conformers increase with increase of the length of the sequence.

**Figure 5.**
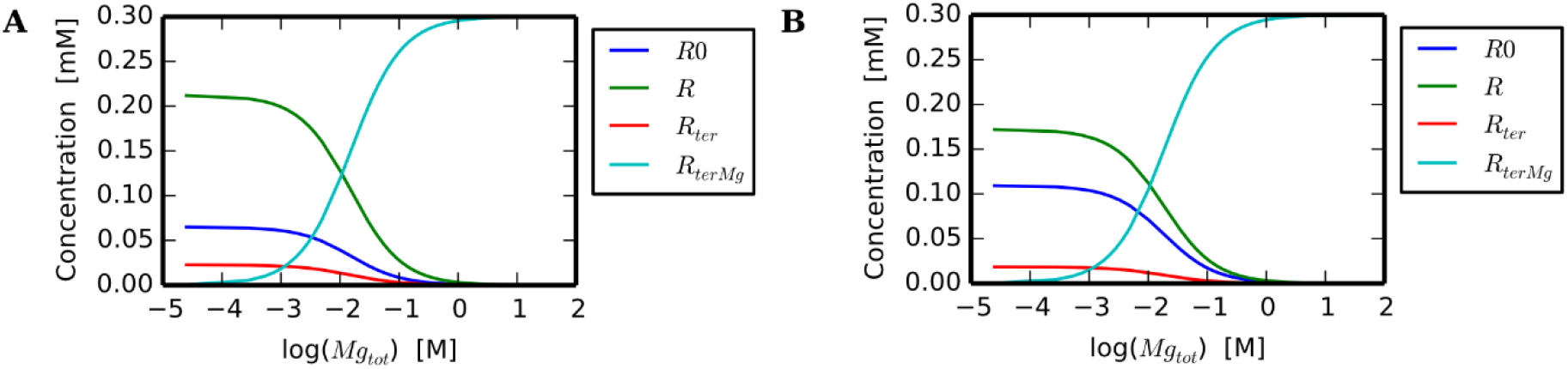
Concentrations of the various RNA species used in *Model 1* as a function of the total magnesium concentration for the wild sequence of lengths 52 and 56 nucleotides. The logarithmic plots show the concentrations of various RNA populations used in *Model 1* including *R*_0_, *R* (the sum of the latter two represent *R*′), *R*_*ter*_ and *R*_*terMg*_. The RNA concentrations are plotted as a function of the total magnesium concentration (*Mg*_*tot*_). The simulations were computed with a total RNA concentration of 0.3 mM for transcript lengths 52 (panel **A**) and 56 (panel **B**) nucleotides, for the wild sequence.

To further illustrate the usefulness of the model, the role of the stability of the P1 helix was studied. In addition to the wild sequence, a set of mutants were chosen (showed in **Figure 6A**), for which titration curves were computed at two different transcription lengths (transcription lengths of 52 and 56). The longer transcription length incorporated the start codon and hence resemble the sequence in bacterial cells. The fraction of binding competent conformers for the different mutants was calculated based on a fixed *K*_*conv*2_ and *K*_*Mg*_, as described in the methods section. The rationale for fixing *K*_*conv*2_ and *K*_*Mg*_ for these mutants is that the mutations in the sequences studied here are not directly involved in the formation of the pseudoknot (hence, *K*_*conv*2_ is unchanged) nor they are located near SAM binding site where the Mg^2+^ is likely to bind (and hence, *K*_*Mg*_ is unlikely to change).

**Figure 6.**
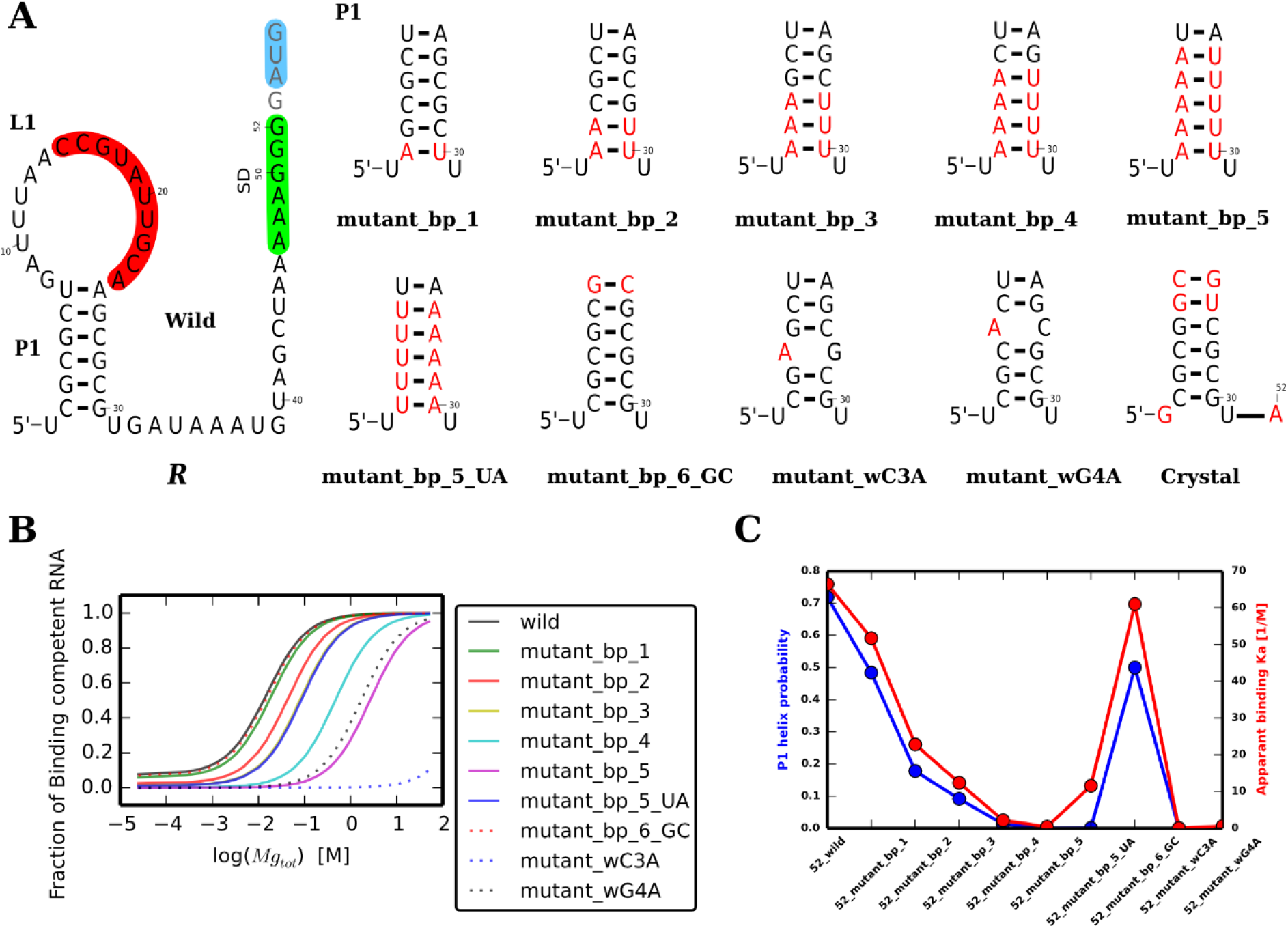
The impact of the stability of the SAM-II riboswitch P1 helix on the fraction of binding competent RNA, for transcript lengths of 52 nucleotides. (**A**) The figure illustrates the wild sequence and its dominating secondary structure, along with the sequences of the P1 helix of the studied mutants. The SD region, the anti-SD and start codon are shaded in green, red and blue, respectively. Nucleotides in grey fonts are those that were truncated in the sequences of length 52 nucleotides, including the experimental and the crystallized sequences, but were included in this study for the wild sequence with transcript length of 56 nucleotides. Point mutations in the studied sequences are depicted in red. (**B**) The plot shows the computed binding competent fractions of the wild sequence along with its various mutants for transcript lengths of 52 nucleotides. (**C**) The plot shows the probability of P1 helix and the apparent binding association constant of Mg^2+^ for the same mutants studied (which are shown in panel **A** and their corresponding titration curves are shown in panel **B**). The corresponding plots illustrated in panels **B** and **C** for transcript lengths of 56 nucleotides are in shown in **Figure S7** (panels **A** and **B**, respectively).

The titration curves for the wild and mutants sequences for transcription lengths 52 and 56, shown in **Figure 6B** and **Figure S7A**, clearly indicate that mutations that destabilize the P1 helix, such as those in the middle of the helix, shift titration curves to the right reflecting that they strongly increase the apparent Mg^2+^ binding dissociation constant (apparent *K*_*d*_) compared to those mutations that have less impact on the stability of the helix. Strikingly, the apparent magnesium binding constant (apparent *K*_*a*_), showed high correlation with probability of the full P1 helix (**Figure 6C** and **Figure S7B**, for the transcript lengths 52 and 56, respectively), even though as described in the methods sections, the chosen secondary structure criteria did not impose the presence of the P1 helix. This indicates that the SAM-II riboswitch is designed in such a way that the P1 helix plays a crucial role in the riboswitch function by providing structural support, and even if it is not directly involved in ligand binding (as it is the case for this riboswitch), it exerts its function by blocking the population of alternative structures.

## 4 DISCUSSION

### 4.1 The treatment of ion-induced RNA conformational changes: non-specific versus specifc bindng

The topic of ion-induced RNA structural changes, particularly those induced by the presence of magnesium ions, has been extensively discussed in the literature [1,34,35]. In general, two classes of magnesium ions are found to bind to RNA. The first class of magnesium ions are those specifically bound to certain sites or residues. Site-specific magnesium ions are often identified from crystal structures, as seen for instance, in the SAM-I [2,36] and SAM-V [11] riboswitches among many other examples in the literature. The second class of ions are those that are non-specifically bound. The latter are found within the solvent layer around the RNA near the backbone phosphate groups, screening the coulomb repulsion between these groups [1]. Several general theories of electrolyte have been used to predict the distribution of ions around nucleic acids, including the Poisson-Boltzmann equation [37–47] and Manning condensation theory [48]. Modified versions of the former to account for various limitations of the original theories have been developed, for instance [49]. Methods have been developed to enable the description of RNA structural changes resulting from the interaction of the solvent ions with the RNA [35,50–52]. Recently, a generalized model of the Manning condensation theory was proposed [53], and have been illustrated in several RNA folding studies [17,20,54].

Hence, often ion binding to nucleic acid, including Mg^2+^, had been treated in terms of non-specific binding, which is thought to improve the electrostatic stability of the RNA, and to a less extent assigned a specific stoichiometry nor a specific binding conformer. Often, when specific binding is assumed, the data is fitted using the Hill equation [55,56], which occasionally may yield non-integer number of bound ions (i.e. fractions of unity), complicating the interpretation of experimental data. A major problem in the treatment of ions-RNA binding is that both site-specific and non-specific ion binding may play a role in the observed overall binding affinities and may contribute to structural changes [1].

In the context of Mg^2+^ induced RNA structural changes; it is important to note that site-specific binding does not exclude the impact of the non-specific ions in the solvation layer. The specific binding of Mg^2+^ ions here is distinct from the non-specific binding in that the former may recognize certain conformers than others, and hence can be understood as a conformational selection mechanism. Whereas the non-specific ions, however, will likely to bind to all RNA molecules in the solution regardless of its structure or folding state, and whether if the difference in the number of ions bound to the compact state compared to the less compact one is sufficient to explain such tertiary interactions had been an on going debate (reviewed in [57]). Indeed, several cases had shown that the change in the charge density between the folded and unfolded state is not enough to provide an explanation for difference in the number of bound magnesium ions between the two states [58,59], and hence cannot justify for the structural changes specifically observed in the presence of Mg^2+^ ions.

### 4.2 Magnesium induced tertiary interactions in riboswitches

Several studies have described the impact of magnesium ions on various riboswitches [2,5,6,20,60,61] including the SAM-II riboswitch [3,7,17,19]. These studies have concluded that magnesium ions aid in stabilizing certain tertiary interactions (e.g. a pseudoknot) that are important for the riboswitch function. The crystal structure of the SAM-I riboswitch, for instance, shows two Mg^2+^ ions which are located at specific sites that aid the formation of RNA tertiary interactions [2,36] and the role of magnesium ions in the formation of the pseudoknot was confirmed by FRET studies [5].

In the specific case of the SAM-II riboswitch presented in this study, previous molecular dynamics simulations and experimental findings proposed a conformational capture mechanism in which conformers that have a nucleated pseudoknot are the binding competent conformers [3,7,18,19]. In other words, SAM recruits the structures that already have a nucleated pseudoknot. The fraction of conformers having this pseudoknot is increased in the presence of magnesium ions in the environment [3,17]. However, NMR experiments showed that this nucleation occurs, with a low probability, even in the absence of both Mg^2+^ and SAM. This observation lead to the conclusion that Mg^2+^ stabilizes that scarcely present state with a nucleated pseudoknot [3]. Therefore, it is reasonable to hypothesize that Mg^2+^ ions favourably bind with the conformers with the already nucleated pseudoknot leading to the increase of its stability. The latter conformers are themselves SAM binding competent conformers. To be able to study the impact of magnesium concentration on riboswitches, a mechanistic model was needed to quantitively describe the structural transitions the riboswitch undergoes upon binding with magnesium ions. Since the SAM-II riboswitch is a translational riboswitch, it is expected that it is thermodynamically driven, and hence equilibrium models, are likely to be able to explain the riboswitch function.

### 4.3 Hints from the structural comparisons between the SAM-II and the SAM-V riboswitches

An important example that needs careful consideration when studying the SAM-II riboswitch is the SAM-V riboswitch [11,21]. Upon visually (i.e. manually, with the aid of molecular graphics software UCSF Chimera [62]) superimposing the purine bases of SAM in the crystal structures of the latter two, a striking similarity in the overall fold of the RNA backbone as well as the SAM configuration is seen (**Figure S1**). Despite this, a hydrated magnesium ion is found near the carboxyl group of SAM in the SAM-V structure [11] and not in SAM-II [14]. The latter magnesium ion is interacting with three water molecules, an oxygen from the phosphate group of the U10 residue and another phosphate oxygen from the G16 residue. The two residues are on opposite sides and roughly in the middle of the L1 loop, and hence the presence a magnesium ion interacting with the two, acts as a bridge to augment the compaction of the two opposite stands of the L1 loop, and hence contributing to the formation of the triplex. At these two residues a kink in the RNA backbone is observed. A closer look at the corresponding sites in the SAM-II (**Figure S1**), reveals that the G17 and U12 in the SAM-II, may play a similar role to the U10 and G16 in the SAM-V structure, respectively, given that the carboxyl group of SAM in the SAM-II structure is directed towards the opposite side of the loop when compared to the SAM-V. An alternative hypothesis is that the SAM-II crystal structure represents one of the possible compact structures of the riboswitch, in which the pseudoknot can be induced by the presence of SAM only, without Mg^2+^. When present, the binding of the ion would induce further structural changes. These hypotheses are further discussed below taking the model that was obtained by superimposing the structures of SAM-II and SAM-V riboswitches as a guide.

**Figure S1** illustrates representative views of the superimposed structures of the SAM-II and SAM-V riboswitches, obtained by specifically superimposing the crystal structures of the two riboswitches (SAM-II and SAM-V), as described above. The resulting model show a striking overall similarity between two riboswitches, as shown in **Figure S1A**. Interestingly, U10 and U12 almost overlap (**Figure S1B**), whereas, G16 and G17 are in proximity, though their corresponding bases are directed differently. SAM configuration exhibits an overall similarity in both models. The major difference is in the geometry of the carboxyl group. The latter carboxyl group in the SAM-V is directed toward the U10 while keeping its proximity to G16. In the SAM-II riboswitch however, the carboxyl group is directed towards the G17 while keeping its proximity to U12. This is also illustrated in panels (**C** to **F** of the same figure).

The wire representation in **Figure S1C** shows a kink in the backbone at G16 (depicted in blue) of the SAM-V and to a less extent at G17 (depicted in red) of the SAM-II riboswitch. Thus, the latter may have been stabilized in the crystal by other means than Mg^2+^, (e.g. crystal packing) or possibly by the exclusive binding of SAM. This sheds the light on the notion that in addition to the major pathway which occurs in the presence of Mg^2+^, the interaction between the two strands of the loop is also attainable through a minor pathway in the absence of Mg^2+^ through exclusive binding of SAM. It can be observed from **Figure S1D** that the distance between the phosphate oxygen of U10 and that of G16 (which are interacting with the magnesium ion) is ~ 5.1 Å, while the distance of the phosphate oxygen of U12 and that of G17 is ~ 7.5 Å (reported here is the shortest distance found between any of the phosphate oxygens of the two residues). The longer distance, which is also reflected in the increased width of the loop and the less prominent kinks in the backbone, is likely due to the absence of the Mg^2+^ ion which bridges the two strands of the loop. Again, it may be that the interaction between the two strands of the loop have been augmented by other forces in the crystal. Even though, in all cases, and despite the likelihood of being less stable compared to the magnesium-bound state, the magnesium free state may still exist in solution and hence proposes an alternative plausible pathway of SAM binding. Taken together, SAM is likely to bind to all conformers having a fully or partially formed pseudoknot, whether it is bound or unbound to magnesium, though the magnesium-bound conformers are likely to be more stable.

The similarity of the Mg^2+^ binding site between SAM-II and SAM-V riboswitches, as illustrated by the superimposed structures shown in panels **E** and **F** of **Figure S1**, respectively, and given the direction of the carboxyl group oxygens of SAM, suggest that residue G17 and U12 in the SAM-II riboswitch may play a similar role to that of the U10 and G16 in the SAM-V, respectively. Another possibility is that Mg^2+^ would induce a structural change that would lead to the re-direction of SAM carboxyl group and the repositioning of G17, resulting in the decrease of the distance between the two strands of the loop. In the latter scenario, the structure of magnesium-bound SAM-II riboswitch would eventually resemble the structure seen in the SAM-V riboswitch. These comparisons therefore propose a hypothesis for the dynamics that may be observed in the presence of the two ligands.

In summary, these structural similarities between the SAM-II and SAM-V along with other experimental conclusions, suggest a site-specific Mg^2+^ binding (as opposed to non-specific binding), through a typical conformational capture mechanism, in which the divalent metal ions specifically interacts with the conformers having a nucleated pseudoknot with a binding energy that is sufficient to stabilize the conformer, and hence increasing the fraction of the riboswitch RNA that can bind with SAM. Similar to the SAM-V, it is likely that Mg^2+^ would also act as a bridge to stabilize interaction between the two strands of the L1 loop. The fact that SAM in the SAM-V structure is stabilized by Mg^2+^ suggests the same with for the SAM-II riboswitch. Indeed, it has been reported that Circular dichroism data confirmed that Mg^2+^ improves SAM binding affinity [7,10]. The proposed model in this study is consistent with such finding in that the presence of Mg^2+^ stabilizes the same conformers that binds with SAM and may stabilizes SAM itself though direct electrostatic interaction, and hence their cooperativity.

### 4.4 Comparison with previous theoretical models and simulations

Several theoretical studies had been conducted on the SAM-II riboswitch folding [10,17,63]. The first quantitative model that addressed magnesium-induced structural changes in the riboswitch attempted to explain the circular dichroism (CD) data and diffusion coefficient [10] though models that propose non-specific magnesium binding. Despite the rigorous treatment, and due to the nature of the CD data, the models are described by a large number of parameters that impedes unique solutions. In a subsequent study, simulations that were based on a generalized Manning condensation theory [53] showed that multiple Mg^2+^ generally interact with the RNA, most of which are directed towards the screening of the destabilizing coulombic potential between the phosphate groups of the backbone [17]. Since these interactions occur with phosphate groups of all nucleotides, such interactions are considered to be non-specific. It is likely that these interactions are present in the presence and absence of the pseudoknot, and it is the difference in the concentration of Mg^2+^ with respect to RNA concentration, is what in principle, should derive the conformational change.

Despite the valuable insights obtained from the latter simulations, which were published before the crystal structure of the SAM-V riboswitch was determined (**Figure S1**), they cannot alone, in this particular case, fully explain the experimental observations. Specifically, the total Mg^2+^ to RNA ratio used in the NMR experiments at 2 mM magnesium, is ~ 2.0 : 0.3 mM i.e. a ratio of ~ 6.67, whereas in the simulations, a much higher relative ratio was used, in which a total Mg^2+^ to RNA ratio of ~ 2.0 : 0.0394 mM (the RNA concentration here is obtained from the volume of the simulation box) was used to obtain a transition, [*Mg*_*tot*_½], similar to that have been obtained from experiments, which was ~ 6 mM. Although provides substantial molecular insights, atomic simulations in general do not take the concentration of the macromolecule into account, which may limit the reproducibility of the experimental conditions in certain bimolecular reactions. Moreover, the average free energy plots of the simulations showed that the presence of Mg^2+^ induces a significant transition from the unfolded U state to the more folded state, whereas as shown by NMR data, that the base pairs in the P1 helix are already formed in the absence of Mg^2+^ and SAM [7]. Nonetheless, these studies demonstrated that the non-specific binding of Mg^2+^ is likely to play an important role even in the presence of a specific magnesium binding site and hence the apparent [*Mg*_*tot*_½] is best explained by taking into account both specific and non-specific magnesium binding.

### 4.5 Comparison of the results obtained by the proposed model with experimental findings

Here, it is illustrated that classical conformational capture models can be used to describe the impact of these divalent ions on the global structure of the RNA assuming site-specific binding. The actual location of such magnesium ion is still not clear. The crystal structure [14], revealed in a single Mg^2+^ ion at the 5′ terminal of the riboswitch, near the phosphate group. Chen et al, however, proposed that the Mg^2+^ may have been delocalized within the crystal structure [7]. This proposal was based on their NMR data, which showed that the spectra for the U12 residue undergo heavy perturbation with the addition of Mg^2+^ ions, suggesting the possibility that this residue may interact with the ion. As discussed earlier, it is likely that the binding sites will resemble the site of the Mg^2+^ ion found in the SAM-V riboswitch, which is in close proximity with SAM.

The presented model revealed that the CEST NMR data, used to monitor the partially formed pseudoknot, can be explained through a 1:1 RNA to Mg^2+^ binding stoichiometry, by direct comparison with the calculated fraction having a partially formed pseudoknot at different concentrations. By fitting the model parameters, the apparent free energy and the calculated [*Mg*_*tot*_½] reveal in an over all transition of [*Mg*_*tot*_½] ~ 10.0 mM (corresponds to an apparent free energy of 2.61 kcal/mol). This transition is in agreement with the [*Mg*_*tot*_½] determined by SEC, considering that this free energy contribution is from the specific binding of Mg^2+^, which is estimated from *NRTln*([*Mg*_*tot*_½]) and is found to be - 3.0 kcal/mol for *N* = 1. The small discrepancy ΔΔ*G* of ~ 0.39 kcal/mol between the calculated and experimentally determined energies is likely due to the non-specific binding. Since, it is assumed that the non-specific binding is similar in all conformers, it does not affect the calculated quantities and results, other than that the total free energy change is equal to the sum of both specific and non-specific free energy contributions.

More specifically, the discrepancy between the calculated and observed [*Mg*_*tot*_½] is because a portion of the magnesium ions are consumed by the non-specific binding to all the RNA species, leading to the slight shift in the overall equilibrium. Indeed, considering non-integer values for 1 < *N* < 2, specifically *N* = 1.4 (using the in-cell model), results in an optimal numerical fit (data not shown), and hence decreases ΔΔ*G* comparted to integer values. However, as justified in the *Methods* section, ions are countable and the ratios between the bound and unbound populations, which reflect the binding strength, is described by the equilibrium constants. Indeed, when fitting binding data using the conventional and widely used Hill equation [55], it is a common practice to vary the Hill coefficient [64] (which may take non-integer values). However, such coefficient may not necessarily reflect the binding stoichiometry, due to multistep binding which is not accounted for in the Hill model. Therefore, the coefficient is best interpreted as an “interaction coefficient” that is an indicator for cooperativity in the case of multiple ligand binding sites [23]. Hence, in the model described here, such non-integer values of *N* are not physically sensible, and it is likely that the arise of ΔΔ*G* is due to the effect of non-specific binding which is usually described by different formalisms that is not considered in this study.

The model presented in this study, is advantageous over previous models that were used to explain the behavior of the SAM-II riboswitch in several aspects. First, the agreement of the model with experimental data is demonstrated using multiple points at specific concentrations, rather than on an overall value such as the midpoint transition [*Mg*_*tot*_½], which is, in a bimolecular interaction, sensitive to the relative concentrations of the interacting species. Second, the predicted fraction with a pseudoknot is illustrated using the total concentrations of both magnesium and RNA that were used in the NMR experiment. Third, the model is consistent with the crystal structure of SAM-V riboswitch which has a very similar structure to the riboswitch used in this study. The model is therefore potentially applicable to the SAM-V as well as other riboswitches that have site-specific bound metal ions. Moreover, the model is not inconsistent with the classical RNA thermodynamics that predicts that the P1 helix probability is stable and exists with high probability even in the absence of Mg^2+^ (**Figure 6C**). This was confirmed by NMR studies [7], in which the authors concluded that most of the P1 base pairs are already formed in the absence of both Mg^2+^ and SAM. Based on a multistep transition, the model overcome limitations of the classical two state transitions, that had been used to explain magnesium induced RNA structural changes [50,55]. Further, unlike several analysis methods used in the literature [3,65], equilibrium constants and standard free energies extracted here are based on a bimolecular interaction and not a unimolecular transition. As a result, the free energies and equilibrium constants therefore do not vary with concentrations. Finally, the model does not exclude, but in fact consistent with, the fact that non-specific binding does contribute to the experimentally observed apparent transition, as shown by the small difference between the calculated and experimental values of [*Mg*_*tot*_½].

Despite these advantages, the model does suffer from several limitations. One limitation is that the model energetic parameters are not based on atomic coordinates, which provides much more details about the system. This means that there is a need for experimental data to be able to estimate the parameters with reasonable confidence. Though this data was available for the case of the SAM-II riboswitch, it may not be necessarily available for other riboswitches. Another limitation of the method is that it does not treat non-specific binding. The assumption here is that magnesium binds similarly to the various RNA species. Such limitations are subjects for future developments.

### 4.6 The integration of ligand concentration, tertiary interaction energetics and RNA folding algorithms extends the range of applications of the model

RNA base-pairing and folding energetics is well established [66–68]. Combining equilibrium analytical models to RNA folding energetics had been extensively used, for example, in oligonucleotide-nucleic acid binding [69]. Further, coupling RNA folding to (small molecule) ligand binding had proven to be a promising method to determine the RNA folding energy landscape in the presence of the cognate ligand [25,70,71]. Here, by incorporating the ligand concentration and tertiary interaction energetics to classical secondary structure prediction, the presented method, therefore, extends the previously reported method [25] to enable predicting the impact of variable concentrations of a ligand on the RNA ensemble, including the nucleation of tertiary interactions using a combination of RNA folding energetics and analytical models.

The latter development enabled the examination of the impact of magnesium on the wild sequence and on certain mutants that will not affect the model parameters. As a case study, this was used to demonstrate the crucial role of the P1 in stabilizing the overall fold, allowing the pseudoknot to be formed by a sufficient fraction of the population, which is a condition that is likely needed for efficient function. It should be noted that care should be taken when using or adapting this model to other riboswitches, as not all riboswitches will necessarily be as the SAM-II riboswitch, in which the P1 seems to have a structural role as opposed to a binding role. The P1 helix in the SAM-I [2,36] and SAM-III [72], for instance, is directly involved in the cognate ligand binding.

### 4.7 Conclusions

In summary, an equilibrium conformational capture model has been developed to model the impact of site-specific magnesium ions on the SAM-II riboswitch. Titration curves calculated based on the fitted model parameters showed good agreement with the experimental data with a 1:1 Mg^2+^ to RNA stoichiometry and not with a higher stoichiometry. The experimentally fitted parameters suggest that the magnesium induced RNA folding occur in a two-step mechanism, in which the pre-existing and dominating P1 helix containing conformers undergo an energetically unfavourable pseudoknot nucleation step, followed by a an energetically favourable Mg^2+^ binding step, and hence stabilizing this state. The model allowed the study of the folding behaviour and the prediction of the apparent binding of the wild sequence and of mutants, with different sequence lengths. The latter studies showed that the P1 helix of the SAM-II riboswitch is a major determinant of the fraction of conformers that are competent for binding with SAM, even though the helix base pairs themselves are not directly involved in the binding.

Overall, this study presented a method that can be used to study riboswitches and was applied here to the SAM-II riboswitch, paving the way to study the role of SAM binding. Because site-specific magnesium binding to RNA is a widespread phenomenon, it is anticipated that such equilibrium model can be adapted to study of other riboswitches. The method contributes to the continuing efforts to delineate the complex free energy landscape sculped by RNA base pairing and tertiary interactions from the one hand, and the binding of magnesium followed by the cognate ligand from another.

## 5 SUPPORTING INFORMATION

Supporting Information is available online.

## 6 ACKNOWLEDGEMENT

The author would like to thank Prof. Fareed Aboul-Ela (Zewail city of Science and Technology), for his helpful comments and discussions.

## 7 CONFLICT OF INTEREST

None declared.

## Notes

### Competing Interest Statement

The authors have declared no competing interest.

