## Supporting Information for "Modelling the impact of magnesium ions concentration on the folding of the SAM-II riboswitch"

Osama Alaidi

Biocomplexity for Research and Consulting, Cairo, Egypt


### 1 Supporting methods

#### 1.1 Deriving an expression for the concentration of the RNA bound to magnesium ions

To be able to predict titration curves, it is convenient to derive an expression for the fraction bound to $Mg^{2+}$, as a function of total concentrations of the reacting moieties (namely, $Mg_{tot}$ & $R_{tot}$), and the equilibrium constants ($K_{ter}$ & $K_{Mg}$).

By rearranging eq. 3 in the main text which is, $K_{Mg} = \frac{R_{terMg}}{R_{ter} \times Mg^N}$ , we obtain,

$$R_{terMg} = K_{Mg} \, . \, R_{ter} \, . \, Mg^N$$

By substituting for $R_{ter}$ from eq. 5, in the above expression, we get,

$$R_{terMg} = K_{Mg} \, . \, K_{ter} \, . \, R^{`} \, . \, Mg^N = R^{`} \, . \, Mg^N \, . \, \alpha$$

Where, $\alpha = K_{Mg} \, . \, K_{ter}$ (eq. S1)

From the mass conservation law, we have,

$$Mg_{tot} = Mg + N.R_{terMg}$$

$$Mg = Mg_{tot} - N.R_{terMg} \qquad \text{(eq. S2)}$$

and we have,

$$R_{tot} = R^{`} + R_{ter} + R_{terMg}$$

$$R_{tot} = R^{`} + K_{ter}R^{`} + R_{terMg}$$

$$R^{`} = \frac{R_{tot} - R_{terMg}}{1 + K_{ter}} = (R_{tot} - R_{terMg})\beta$$

Where, $\beta = \frac{1}{1+K_{ter}}$ (eq. S3)

Substitution of equations S2 and S3 into S1 gives,

$$R_{terMg} = R^{`} \, . \, Mg^N \, . \, \alpha \qquad \text{(eq. S4)}$$

$$= \left(R_{tot} - R_{terMg}\right)\beta \left(Mg_{tot} - N.R_{terMg}\right)^N \alpha$$

$$= (R_{tot} - R_{terMg}) \left(Mg_{tot} - N.R_{terMg}\right)^N \gamma \qquad \text{(eq. S5)}$$

*Where*, $\gamma = \alpha\beta = \frac{K_{Mg} \, . \, K_{ter}}{1+K_{ter}} = \frac{K_{Mg}}{\frac{1}{K_{ter}} + 1}$

#### 1.1.1 *For the case of* $N = 1$ *(i.e. 1:1 stoichiometry), equation S5 simplifies to,*

$$R_{terMg} = (R_{tot} - R_{terMg})\,(Mg_{tot} - R_{terMg})\,\gamma \qquad \text{(eq. S6)}$$

By dividing both sides of the equation by $\gamma$ and then from each side of the equation subtracting $\frac{R_{terMg}}{\gamma}$, followed by multiplying terms in the brackets, we obtain,

$$(R_{terMg})^2 - R_{terMg}Mg_{tot} - R_{terMg}R_{tot} - \frac{R_{terMg}}{\gamma} + R_{tot}Mg_{tot} = 0$$

$$(R_{terMg})^2 + \left(-Mg_{tot} - R_{tot} - \frac{1}{\gamma}\right)R_{terMg} + R_{tot}Mg_{tot} = 0$$

The solution to the quadratic, is given by,

$$R_{terMg} = \frac{-b \pm \sqrt{b^2 - 4ac}}{2a}$$

Where,

$a = 1,$ $\quad b = \left(-Mg_{tot} - R_{tot} - \frac{1}{\gamma}\right)$ $\quad$ and $\quad c = R_{tot}Mg_{tot}$

The quadratic equation has two possible solutions for $R_{terMg}$ , which are given by the expression,

$$R_{terMg} = \frac{Mg_{tot} + R_{tot} + \frac{1}{\gamma} \pm \sqrt{(-Mg_{tot} - R_{tot} - \frac{1}{\gamma})^2 - 4R_{tot}Mg_{tot}}}{2}$$

and $\quad \gamma = \frac{K_{Mg}}{\frac{1}{K_{ter}} + 1}$ $\qquad$ (eq. S7)

Only one of the two roots is physically sensible, and can represent concentrations (i.e. positive real number, with a maximum value equal or less than $R_{tot}$).

#### 1.1.2 *For the case of* $N = 2$ *(i.e. 2* $Mg^{2+}$ *: 1 RNA stoichiometry)*

By substituting for $N = 2$, equation S5 becomes,

$$R_{terMg} = (R_{tot} - R_{terMg})\,(Mg_{tot} - 2\,R_{terMg})^2\,\gamma$$

$$= (R_{tot} - R_{terMg})\,(Mg_{tot}^{\;2} - 4\,Mg_{tot}R_{terMg} + 4R_{terMg}^{\;2})\,\gamma$$

$$= (-4R_{terMg}^{\;3} + 4\,R_{tot}R_{terMg}^{\;2} + 4\,Mg_{tot}R_{terMg}^{\;2} - Mg_{tot}^{\;2}R_{terMg} - 4\,R_{tot}Mg_{tot}R_{terMg} + R_{tot}Mg_{tot}^{\;2})\,\gamma$$

By dividing both sides of the equation by $\gamma$, then subtracting $\frac{R_{terMg}}{\gamma}$ from both sides, and collecting similar terms, we obtain,

$$(-4)\, R_{terMg}{}^{3} + (4\, Mg_{tot} + 4\, R_{tot})\, R_{terMg}{}^{2} + (-Mg_{tot}{}^{2} - 4R_{tot}Mg_{tot} - \frac{1}{\gamma})\, R_{terMg} + R_{tot}Mg_{tot}{}^{2} = 0$$

Hence, the cubic equation is in the form

$$a\, R_{terMg}{}^{3} + b\, R_{terMg}{}^{2} + c\, R_{terMg} + d = 0$$

Where,

$a = -4$ , $b = 4Mg_{tot} + 4R_{tot}$ , $c = -Mg_{tot}{}^{2} - 4R_{tot}Mg_{tot} - \frac{1}{\gamma}$ and $d = R_{tot}Mg_{tot}{}^{2}$

The cubic equation has three possible solutions for $R_{terMg}$, only one of which is physically sensible and can represent concentrations (i.e. positive real number, with maximum value equal or less than $R_{tot}$). The solutions (i.e. roots) for each were obtained by 'numpy.roots()' function in the python NumPy library, which returns the roots of a polynomial given its coefficients [1]. The latter uses an algorithm that computes the eigenvalues of the companion matrix [1,2].

#### 1.2 Deriving an expression for the concentration and fraction of the RNA bound to magnesium ions inside the cell

Inside bacterial cells, we can assume that $Mg \cong Mg_{tot} \cong Mg_{cyt}$ , where $Mg_{cyt}$ is the bulk (free) concentration of magnesium in the cytoplasm.

Hence from equation S4, $R_{terMg}$ is equal to,

$$R_{terMg} = R^{`} \,.\, (Mg)^{N} \,.\, \alpha \cong (R_{tot} - R_{terMg})(Mg_{cyt})^{N} \beta\, \alpha$$

(eq. S8)

Let the constant quantity, $\theta = (Mg_{cyt})^{N} \beta\, \alpha$ , then

$$R_{terMg} = \theta(R_{tot} - R_{terMg})$$

Solving for $R_{terMg}$ this gives,

$$R_{terMg} = \frac{R_{tot}}{\frac{1}{\theta} + 1}$$

Where, $\theta = \frac{K_{Mg} \times (Mg_{cyt})^{N}}{\frac{1}{K_{ter}} + 1}$ and $N$ is the number of moles of $Mg^{2+}$ ions that binds to 1 mole of RNA.

The assumption simplifies the estimation of the concentration and fraction of RNA bound to magnesium for any given stoichiometry.

### 2 Supporting Figures

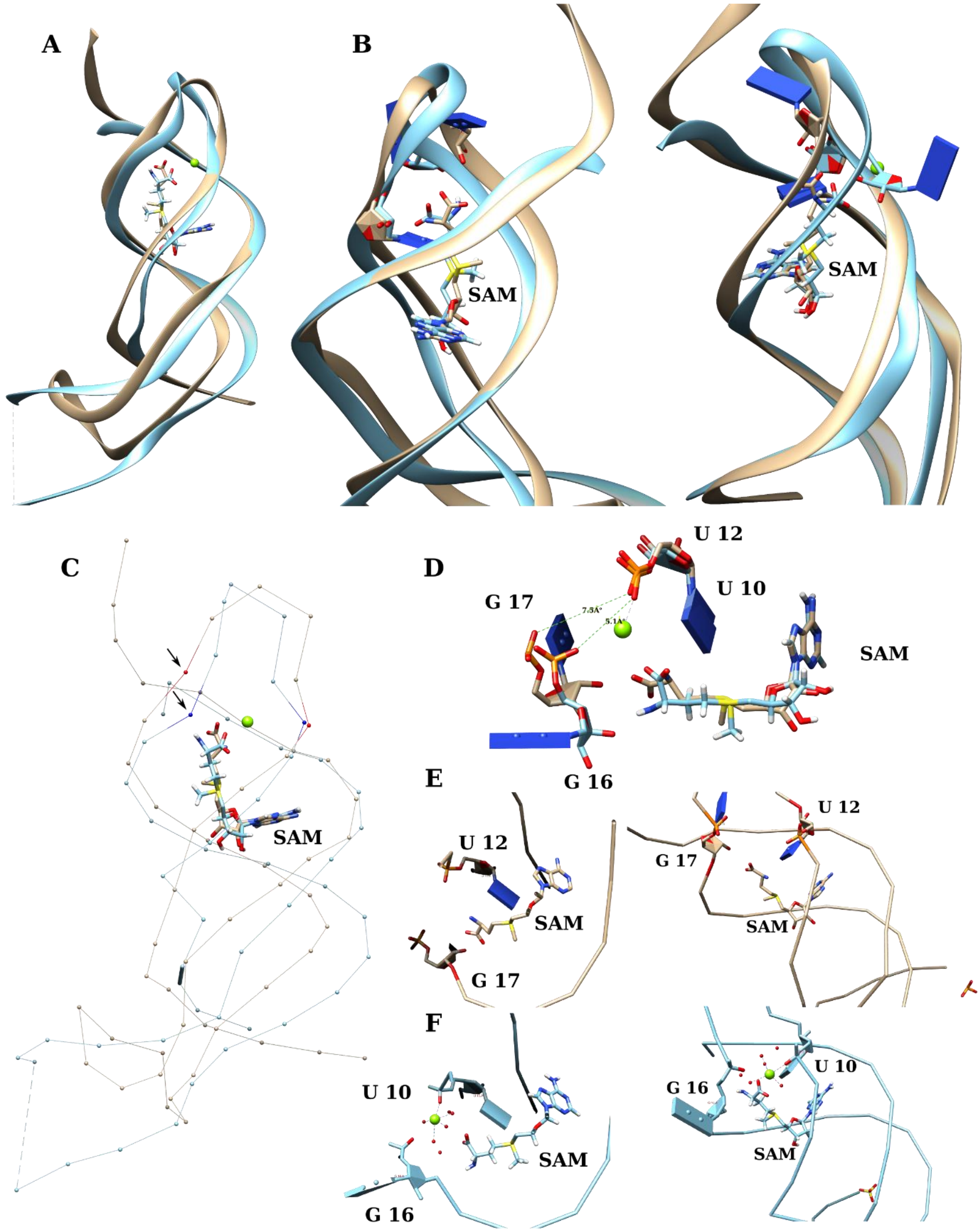


**Figure S1. Structural similarity of the SAM-II and SAM-V riboswitches as revealed by superimposing their crystal structures.**

Representative views of the superimposed structures of the SAM-II and SAM-V riboswitches (PDB ID *2qwy* and *6fz0*, respectively). In all panels the ribbon and wire representations are depicted in gold and cyan, for the SAM-II and the SAM-V riboswitches, respectively. Oxygens of the SAM carboxyl group and that of the phosphate groups of nucleotides of interest are depicted in red. (**A**) Backbone in ribbon representation showing the overall similarity of the two structures. (**B**) Closeup front and back views of **A**, with residues (U10 and G16) from the SAM-V and (U12 and G17) from the SAM-II riboswitches are shown. (**C**) Wire representations showing a kink (marked by arrows) in the RNA backbone at G16 of the SAM-V riboswitch and to a less extent a similar kink can be observed at G17 of the SAM-II riboswitch (residues U10 and G16 are depicted in blue, whereas U12 and G17 are depicted in red). (**D**) The two pairs of residues (U10, G16 and U12, G17) of the superimposed structures are shown along with SAM. Panels (**E**) and (**F**) illustrate the similarity of the $Mg^{2+}$ binding site along with the magnesium-bound water molecules, via two different views of the superimposed structures, showing only the SAM-II in panel **E** and it corresponding views of the SAM-V in panel **F**. The Magnesium ion and bound water molecules are in green and red colors, respectively.

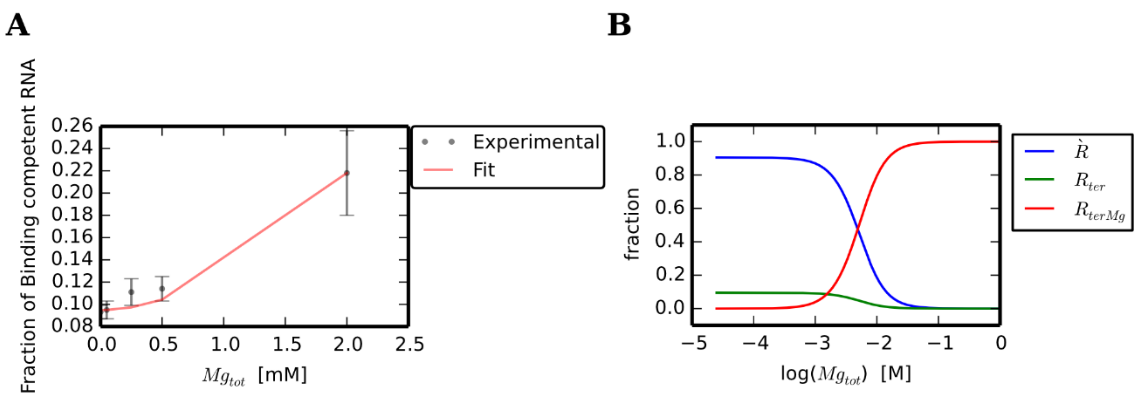


**Figure S2. Fractions of various RNA species as a function of magnesium concentration assuming a 1:2 RNA to $Mg^{2+}$ stoichiometry.**

Panel (**A**) shows a plot for the experimental data (and their associated error bars) versus the calculated fraction of binding competent conformers, using $N = 2$. The calculated fraction is obtained by the global fitting of model parameters to the fraction of RNA with a nucleated pseudoknot inferred from NMR CEST experimental data and reported by Chen et al [3], in which the base paring of G22 and C43 was monitored. Panel (**B**) shows a plot of the calculated fraction of each RNA population/species at an increasing magnesium concentration calculated from the globally fitted parameters for $N = 2$. The fractions of each of $R^{`}$, $R_{ter}$ and $R_{terMg}$ (fraction bound) as a function of magnesium concentration (on a logarithmic scale) are shown in blue, green and red, respectively. From structural point of view, $R^{`}$ and fraction binding competent ($R_{ter} + R_{terMg}$) represent the fractions of RNA without and with a nucleated pseudoknot, respectively.

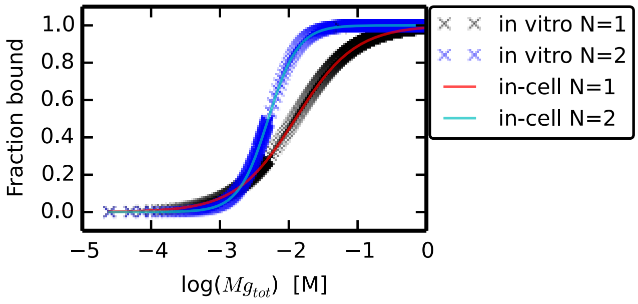


**Figure S3. The *fraction of bound* RNA calculated with various model assumptions.**

The figure shows the *fraction bound* i.e. ($R_{terMg}$), plotted as a function of the total magnesium molar concentration for $N = 1$ and $N = 2$, each for both in vitro and in-cell approximations. The logarithmic plot enables the comparison between the probability of the RNA bound to $Mg^{2+}$ ions as predicted by the various assumptions.

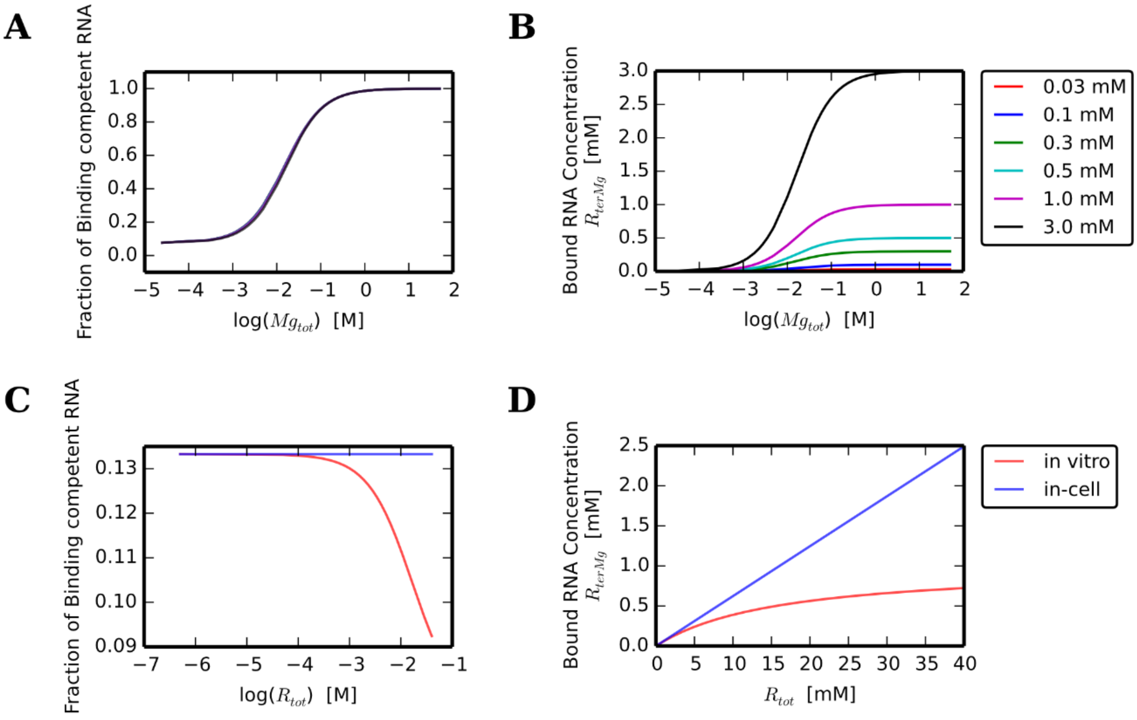


**Figure S4. The effect of varying the total RNA concentration ($R_{tot}$).**

The figure shows representative plots that illustrate the effect of varying the total RNA concentration ($R_{tot}$) on the concentrations of the bound RNA and on the fractions of the binding competent RNA, using both in vitro and in-cell models. Panels (**A** and **B**) illustrate the fractions of binding competent RNA and the bound RNA concentrations, respectively, as a function of magnesium concentration, calculated for various $R_{tot}$ using the in vitro model. Panels (**C** and **D**) show the same quantities on the y-axis as in **A** and **B**, respectively, but for a fixed total magnesium concentration of 1 mM calculated by varying $R_{tot}$ instead, with the former shown on a logarithmic scale. For comparison, the latter plots (in **C** and **D**) are shown for both the in vitro (red line) and the in-cell (blue line) models.

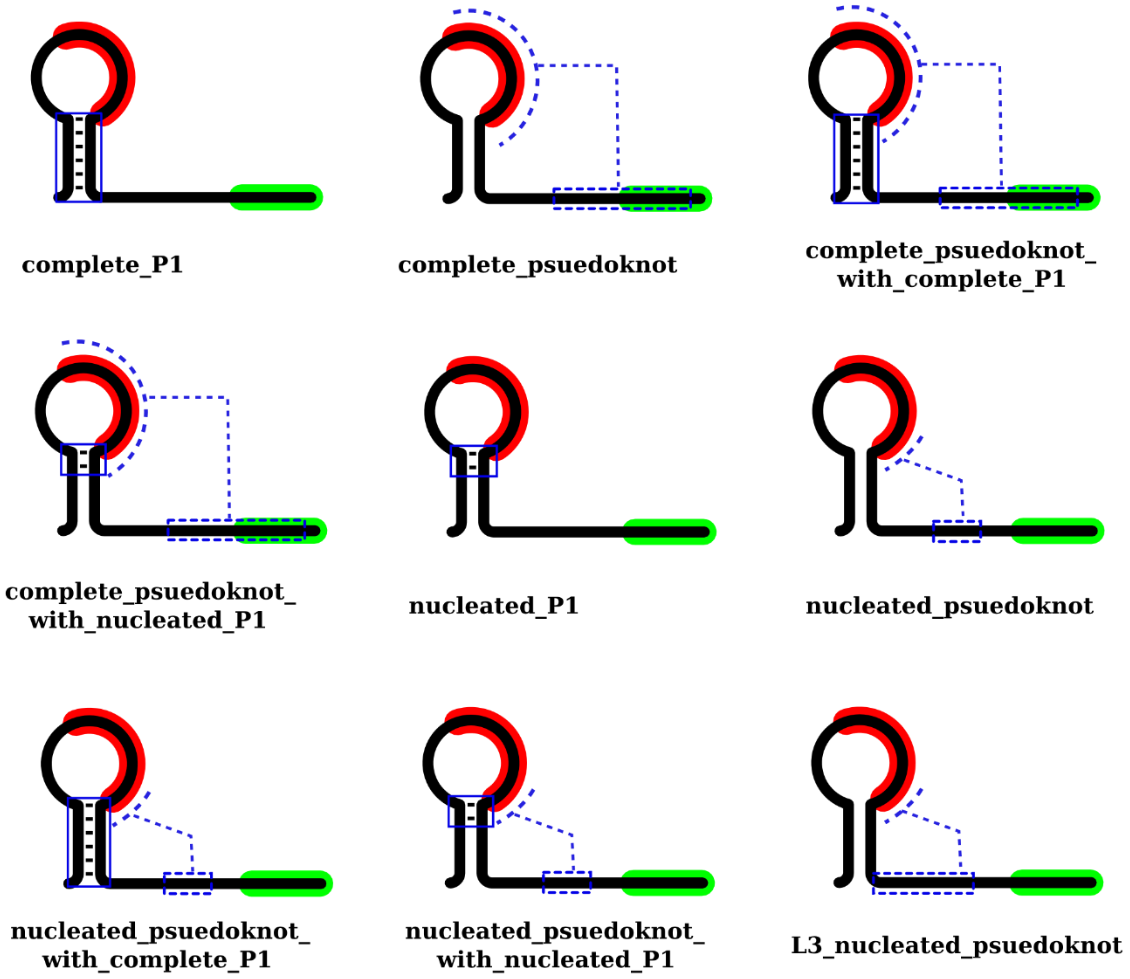


**Figure S5. Secondary Structure conditions used to define the species $R$.**

The figure illustrates the different secondary structure conditions examined. These conditions were used to define conformers forming species $R$ (sequences and positions are listed in **Table S5**). As shown in the skeleton diagrams, the P1 helix nucleotides that must be base paired to each other in order to satisfy a certain condition, are connected by black lines and the part of the helix examined is boxed (blue solid lines). Structural motifs that cannot be predicted by classical RNA secondary structure algorithms, in which certain nucleotides need to be unpaired to form a proposed tertiary interaction, are marked or boxed with dashed blue lines.

**A**

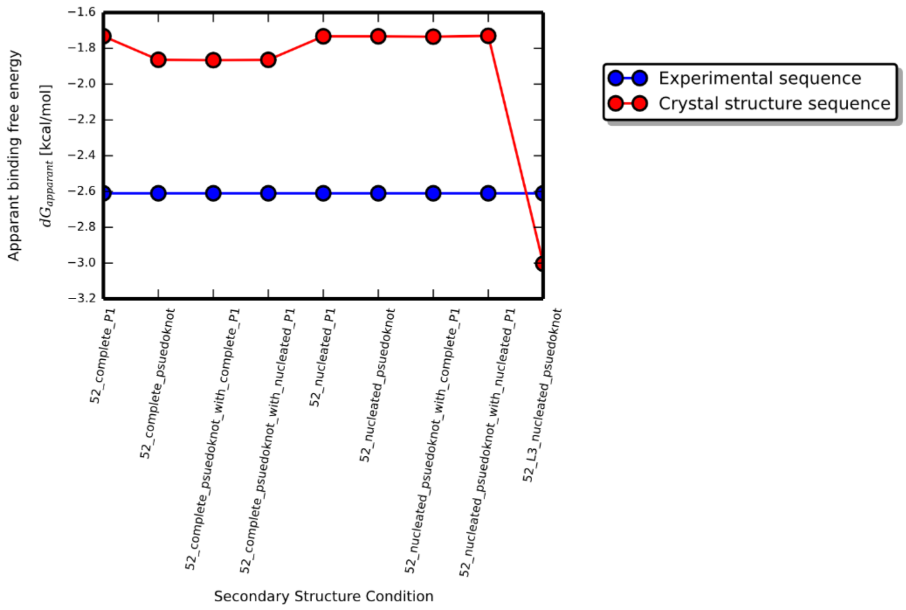


**B**

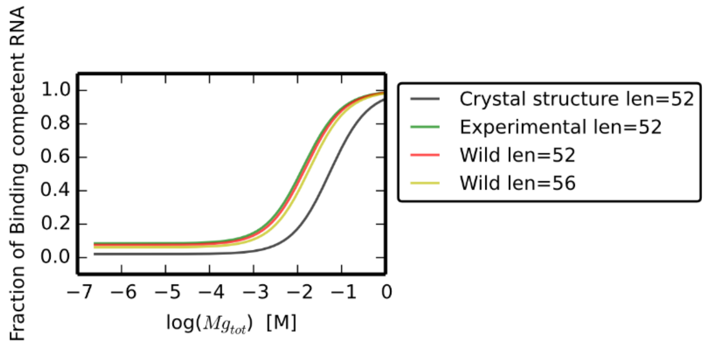


**Figure S6. Examination of the various assumptions made about the secondary structure conditions used to define the species $R$ and their impact on the apparent $Mg^{2+}$ binding free energy.**

(**A**) The plot shows the apparent binding free energies of the crystal structure sequence along with the experimental sequence calculated using various assumptions about the minimal secondary structure conditions required for the pseudoknot formation. The prefix 52 refer to the transcription length and the label on the x-axis specifies the condition used to define the population $R$ in *Model 1*. The analysis does not confirm a specific condition nor a conversion pathway. However, the analysis can be used to reject certain conditions based on its consistency with experimental trends. (**B**) The plot shows the fraction of binding competent conformers, at an increasing magnesium concentration, for the crystal structure, experimental, wild (length= 52 nucleotides) and wild (length = 56 nucleotides) sequences. The titration curves were calculated assuming that the conformers $R$, are those having a secondary structure with a *nucleated pseudoknot* as defined in **Table S5** and schematically shown in **Figure S5**.

**A** **B**

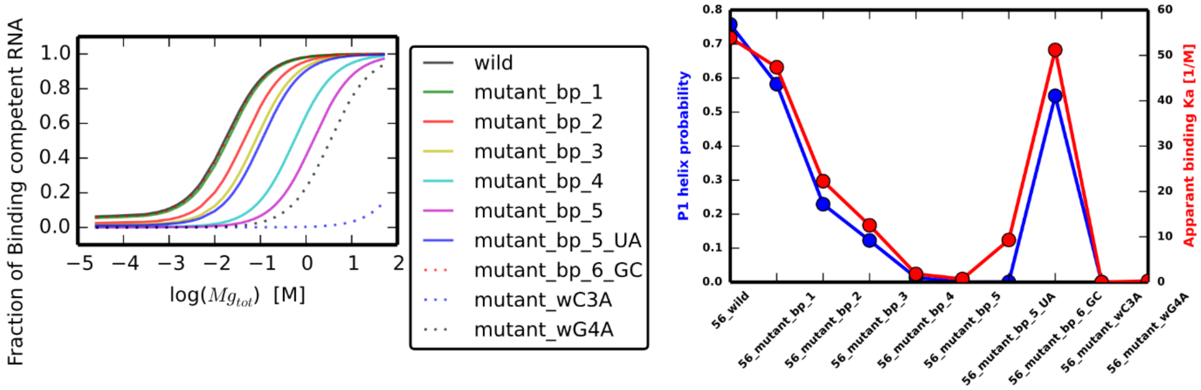


**Figure S7. The impact of the stability of the SAM-II riboswitch P1 helix on the fraction of binding competent conformers, for transcript lengths of 56 nucleotides.**

(**A**) The plot shows the computed binding competent fractions of the wild sequence along with its various mutants for transcript lengths of 56 nucleotides. (**B**) The plot shows the probability of P1 helix and the apparent binding association constant of $Mg^{2+}$ for the same mutant studied in panel **A** (which are schematically shown in **Figure 6A**). The corresponding plots illustrated in panels **A** and **B** for transcript lengths of 52 nucleotides are shown in **Figure 6** (panels **B** and **C,** respectively).

### 3 Supporting Tables

| RNA species Symbol | Concentrations (mM) | Probability % |
|---|---|---|
| $R^{`}$ | 27.21E-2 | 90.7 |
| $R_{ter}$ | 2.79E-2 | 9.3 |
| $R_{terMg}$ | 0.00 | 0.0 |
| $R_{tot}$ | 0.3 | 100 |
| **Model parameters** | | |
| $K_{ter}$ | 102.54E-3 | |
| $\Delta G_{ter}$ | 1.35 kcal/mol | |

**Table S1. The concentrations and probabilities of the various RNA populations in the absence of magnesium ions.**

The table shows the concentrations and probabilities of the various RNA species in the absence of $Mg^{2+}$ ions at a 0.3 mM RNA concentration. The values are based on the measurements reported in [3]. The concentrations (or equally probabilities) were used to calculate the association constant $K_{ter}$. As described in the text, $R^{`}$ is RNA population without a pseudoknot, $R_{ter}$ is RNA population with a pseudoknot that are *not* bound to $Mg^{2+}$, and $R_{terMg}$ is RNA population with a pseudoknot *and* are bound to $Mg^{2+}$ ions.

| 0.05 mM $Mg^{2+}$ | | |
|---|---|---|
| **RNA/ Magnesium species Symbol** | **Concentrations (mM)** | **Probability %** |
| $R^{`}$ | 2.72E-01 | 90.50 |
| $R_{ter}$ | 2.78E-02 | 9.28 |
| $R_{terMg}$ | 6.62E-04 | 0.22 |
| $R_{tot}$ | 0.3 | 100.00 |
| $Mg$ | 4.93E-02 | |
| $R_{terMg}$ | 6.62E-04 | |
| $Mg_{tot}$ | 0.05 | |
| **Model parameters** | | |
| $K_{Mg}$ | 481.63 $M^{-1}$ | |
| $Kd_{Mg}$ | 2.08E-03 M | |
| $\Delta G_{Mg}$ | -3.66 kcal/Mol | |

| 0.25 mM $Mg^{2+}$ | | |
|---|---|---|
| **RNA/ Magnesium species Symbol** | **Concentrations (mM)** | **Probability %** |
| $R^{`}$ | 2.67E-01 | 88.90 |
| $R_{ter}$ | 2.73E-02 | 9.12 |
| $R_{terMg}$ | 5.95E-03 | 1.98 |
| $R_{tot}$ | 0.3 | 100.00 |
| $Mg$ | 2.44E-01 | |
| $R_{terMg}$ | 5.95E-03 | |
| $Mg_{tot}$ | 0.25 | |
| **Model parameters** | | |
| $K_{Mg}$ | 892.10 $M^{-1}$ | |
| $Kd_{Mg}$ | 1.12E-03 M | |
| $\Delta G_{Mg}$ | -4.03 kcal/Mol | |

| 0.5 mM $Mg^{2+}$ | | |
|---|---|---|
| **RNA/ Magnesium species Symbol** | **Concentrations (mM)** | **Probability %** |
| $R^{`}$ | 2.66E-01 | 88.60 |
| $R_{ter}$ | 2.73E-02 | 9.08 |
| $R_{terMg}$ | 6.95E-03 | 2.32 |
| $R_{tot}$ | 0.3 | 100.00 |
| $Mg$ | 4.93E-01 | |
| $R_{terMg}$ | 6.95E-03 | |
| $Mg_{tot}$ | 0.5 | |
| **Model parameters** | | |
| $K_{Mg}$ | 516.90 $M^{-1}$ | |
| $Kd_{Mg}$ | 1.94E-03 M | |
| $\Delta G_{Mg}$ | -3.70 kcal/Mol | |

| 2.0 mM $Mg^{2+}$ | | |
|---|---|---|
| **RNA/ Magnesium species Symbol** | **Concentrations (mM)** | **Probability %** |
| $R^`$ | 2.35E-01 | 78.20 |
| $R_{ter}$ | 2.41E-02 | 8.02 |
| $R_{terMg}$ | 4.13E-02 | 13.78 |
| $R_{tot}$ | 0.3 | 100.00 |
| $Mg$ | 1.96 | |
| $R_{terMg}$ | 4.13E-02 | |
| $Mg_{tot}$ | 2.0 | |
| **Model parameters** | | |
| $K_{Mg}$ | 877.53 $M^{-1}$ | |
| $Kd_{Mg}$ | 1.14E-03 M | |
| $\Delta G_{Mg}$ | -4.02 kcal/Mol | |

**Table S2. The concentrations and probabilities of the various RNA populations in the presence of magnesium ions, and their corresponding free and bound magnesium ion concentrations, for 1:1 stoichiometry ($N = 1$).**

The table shows the concentrations and probabilities of the various RNA species in the presence of 0.05, 0.25, 0.5 and 2 mM $Mg^{2+}$, at a 0.3 mM RNA concentration. The concentrations were used to calculate the magnesium ions binding association constant $K_{Mg}$. The concentrations of free and bound $Mg^{2+}$ ions in the same conditions are also listed. The calculations in this table were performed for 1:1 stoichiometry (i.e. with $N = 1$), using single point experimental data reported in [3]. As described in the text, $R^`$ is RNA population without a pseudoknot, $R_{ter}$ is RNA population with a pseudoknot that are *not* bound to $Mg^{2+}$, and $R_{terMg}$ is RNA population with a pseudoknot *and* are bound to $Mg^{2+}$( also used to describe the *Bound* $Mg^{2+}$). $R_{tot}$, $Mg$ and $Mg_{tot}$ are the total RNA, *free* $Mg^{2+}$ and *total* $Mg^{2+}$ concentrations, respectively.

| 0.05 mM $Mg^{2+}$ | | |
|---|---|---|
| **RNA/ Magnesium species Symbol** | **Concentrations (mM)** | **Probability %** |
| $R`$ | 2.72E-01 | 90.50% |
| $R_{ter}$ | 2.78E-02 | 9.28% |
| $R_{terMg}$ | 6.62E-04 | 0.22% |
| $R_{tot}$ | 3.00E-01 | 100.00% |
| $Mg$ | 4.87E-02 | |
| $R_{terMg}$ | 1.32E-03 | |
| $Mg_{tot}$ | 0.05 | |
| **Model parameters** | | |
| $K_{Mg}$ | 1.00E+07 $M^{-1}$ | |
| $Kd_{Mg}$ | 9.97E-08 M | |
| $\Delta G_{Mg}$ | -9.55 kcal/mol | |

| 0.25 mM $Mg^{2+}$ | | |
|---|---|---|
| **RNA/ Magnesium species Symbol** | **Concentrations (mM)** | **Probability %** |
| $R`$ | 2.67E-01 | 88.90% |
| $R_{ter}$ | 2.73E-02 | 9.12% |
| $R_{terMg}$ | 5.95E-03 | 1.98% |
| $R_{tot}$ | 3.00E-01 | 100.00% |
| $Mg$ | 2.38E-01 | |
| $R_{terMg}$ | 1.19E-02 | |
| $Mg_{tot}$ | 0.25 | |
| **Model parameters** | | |
| $K_{Mg}$ | 3.84E+06 $M^{-1}$ | |
| $Kd_{Mg}$ | 2.60E-07 M | |
| $\Delta G_{Mg}$ | -8.98 kcal/mol | |

| 0.5 mM $Mg^{2+}$ | | |
|---|---|---|
| **RNA/ Magnesium species Symbol** | **Concentrations (mM)** | **Probability %** |
| $R`$ | 2.66E-01 | 88.60% |
| $R_{ter}$ | 2.73E-02 | 9.08% |
| $R_{terMg}$ | 6.95E-03 | 2.32% |
| $R_{tot}$ | 3.00E-01 | 100.00% |
| $Mg$ | 4.86E-01 | |
| $R_{terMg}$ | 1.39E-02 | |
| $Mg_{tot}$ | 0.5 | |
| **Model parameters** | | |
| $K_{Mg}$ | 1.08E+06 $M^{-1}$ | |
| $Kd_{Mg}$ | 9.27E-07 M | |
| $\Delta G_{Mg}$ | -8.23 kcal/mol | |

| 2.0 mM $Mg^{2+}$ | | |
|---|---|---|
| **RNA/ Magnesium species Symbol** | **Concentrations (mM)** | **Probability %** |
| $R^{`}$ | 2.35E-01 | 78.20% |
| $R_{ter}$ | 2.41E-02 | 8.02% |
| $R_{terMg}$ | 4.13E-02 | 13.78% |
| $R_{tot}$ | 3.00E-01 | 100.00% |
| $Mg$ | 1.92 | |
| $R_{terMg}$ | 8.27E-02 | |
| $Mg_{tot}$ | 2.0 | |
| **Model parameters** | | |
| $K_{Mg}$ | 4.68E+05 $M^{-1}$ | |
| $Kd_{Mg}$ | 2.14E-06 M | |
| $\Delta G_{Mg}$ | -7.74 kcal/mol | |

**Table S3. The concentrations and probabilities of the various RNA populations in the presence of magnesium ions, and their corresponding free and bound magnesium ion concentrations, for 1:2 RNA to $Mg^{2+}$ stoichiometry ($N = 2$).**

The table shows the concentrations and probabilities of the various RNA species in the presence of 0.05, 0.25, 0.5 and 2 mM $Mg^{2+}$, at a 0.3 mM RNA concentration. The concentrations were used to calculate the magnesium ions binding association constant $K_{Mg}$. The concentrations of free and bound $Mg^{2+}$ ions in the same conditions are also listed. The calculations in this table were performed for 1:2 RNA to $Mg^{2+}$ stoichiometry (i.e. with $N = 2$), using single point experimental data reported in [3]. As described in the text, $R^{`}$ is RNA population without a pseudoknot, $R_{ter}$ is RNA population with a pseudoknot that are *not* bound to $Mg^{2+}$, and $R_{terMg}$ is RNA population with a pseudoknot *and* are bound to $Mg^{2+}$( also used to describe the *Bound* $Mg^{2+}$). $R_{tot}$, $Mg$ and $Mg_{tot}$ are the total RNA, *free* $Mg^{2+}$ and *total* $Mg^{2+}$ concentrations, respectively.

| **Model Parameter (Model 2)** | **Values ($N = 2$) *in vitro*** | **Values ($N = 2$) *In-cell*** |
|---|---|---|
| $K_{ter}$ | 0.105 | 0.105 |
| $Kd_{ter}$ | 9.54 | 9.54 |
| $\Delta G_{ter}$ | 1.34 kcal/Mol | 1.34 kcal/Mol |
| $K_{Mg}$ | $4.51 \times 10^5$ $M^{-1}$ | $4.15 \times 10^5$ $M^{-1}$ |
| $Kd_{Mg}$ | $2.22\times 10^{-6}$ M | $2.41\times 10^{-6}$ M |
| $\Delta G_{Mg}$ | -7.71 kcal/Mol | -7.66 kcal/Mol |
| $[Mg_{tot}½]$ | 4.54 mM | 4.54 mM |
| $\Delta G_{apparent}$ | -6.32 kcal/Mol | -6.31 kcal/Mol |
| $Chi^2$ | $23.47\times 10^{-3}$ | $24.14\times 10^{-3}$ |
| $Chi^2\ err$ | 2.347 | 2.408 |

| **Model Parameter (Model 1)** | **Values ($N = 2$) *in vitro*** | **Values ($N = 2$) *In-cell*** |
|---|---|---|
| $K_{conv1}$ | 22.54 | 22.54 |
| $\Delta G_{conv1}$ | -1.85 kcal/Mol | -1.85 kcal/Mol |
| $K_{conv2}$ | 0.110 | 0.110 |
| $\Delta G_{conv2}$ | 1.31 kcal/Mol | 1.31 kcal/Mol |

**Table S4. Model parameters obtained by global fitting for 1:2 RNA to $Mg^{2+}$ stoichiometry ($N = 2$).**

The table on the left shows the model parameters, i.e. equilibrium constants and corresponding free energies, that were obtained from global fitting to experimental data (*Model 2*), with $N = 2$. The table on the right lists parameters used in *Model 1*, taking the presence of unpaired C23, A24, U40, A41 and G42 as the criteria for the nucleation of the pseudoknot (see **Methods** section for further details). The latter condition in the secondary structures means that contacts involved in the pseudoknot nucleation are accessible, allowing the interaction (i.e. base pairing) between CA (nucleotides 23:24) and UAG (nucleotides 40:42) to occur.

| Condition | Nucleotide position | Sequence | Structure | Closed Helix |
|---|---|---|---|---|
| *52_complete_P1* | 2:7*25:30 | CGCGCU*AGCGCG | ((((((*)))))) | True |
| *052_complete_psuedoknot* | 15:24*40:51 | CCGUAUUGCA*UAGCUAAAAAGG | ..........*............ | N/A |
| *052_complete_psuedoknot_with_complete_P1* | 2:7*15:24*40:51*25:30 | CGCGCU*CCGUAUUGCA*UAGCUAAAAAGG*AGCGCG | ((((((*..........*............*)))))) | True |
| *052_complete_psuedoknot_with_nucleated_P1* | 6:7*15:24*40:51*25:26 | CU*CCGUAUUGCA*UAGCUAAAAAGG*AG | ((*..........*............*)) | True |
| *052_nucleated_P1* | 6:7*25:26 | CU*AG | ((*)) | True |
| *052_nucleated_psuedoknot* | 23:24*40:42 | CA*UAG | ..*... | N/A |
| *052_nucleated_psuedoknot_with_complete_P1* | 2:7*23:24*40:42*25:30 | CGCGCU*CA*UAG*AGCGCG | ((((((*..*...*)))))) | True |
| *052_nucleated_psuedoknot_with_nucleated_P1* | 6:7*23:24*40:42*25:26 | CU*CA*UAG*AG | ((*..*...*)) | True |
| *052_L3_nucleated_psuedoknot* | 23:24*31:42 | CA*UGAUAAAUGUAG | ..*............ | N/A |

**Table S5. Secondary structure conditions that were used for defining conformers constituting species $\boldsymbol{R}$.**

The table lists the nucleotide positions, sequences, and secondary structures (in bracket notation) of the conditions that were examined and used for defining conformers constituting species $R$. The symbol “*” in the table refers to any number or any type of nucleotides, while “N/A” refers to “not applicable”. For example, for the condition of *complete_P1 (i.e.* conformers that have a complete P1 helix*)*, conformers were considered to belong to the species $R$ if their nucleotides at positions 2 to 7 and 25 to 30, whose corresponding sequences are CGCGCU and AGCGCG, respectively, were found to have secondary structures of (((((( and )))))) for the latter two regions, respectively, with any sequence/structure in between, provided that they formed a closed helix (i.e. closed helix condition was ”True”).
